# Data integration uncovers the metabolic bases of phenotypic variation in yeast

**DOI:** 10.1101/2020.06.23.166405

**Authors:** Marianyela Petrizzelli, Dominique de Vienne, Thibault Nidelet, Camille Noûs, Christine Dillmann

## Abstract

The relationship between different levels of integration is a key feature for understanding the genotype-phenotype map.

Here, we describe a novel method of integrated data analysis that incorporates protein abundance data into constraint-based modeling to elucidate the biological mechanisms underlying phenotypic variation. Specifically, we studied yeast genetic diversity at three levels of phenotypic complexity in a population of yeast obtained by pairwise crosses of eleven strains belonging to two species, *Saccharomyces cerevisiae* and *S. uvarum*. The data included protein abundances, integrated traits (life-history/fermentation) and computational estimates of metabolic fluxes.

Results highlighted that the negative correlation between production traits such as population carrying capacity (*K*) and traits associated with growth and fermentation rates (*J*_max_) is explained by a differential usage of energy production pathways: a high *K* was associated with high TCA fluxes, while a high *J*_max_ was associated with high glycolytic fluxes. Enrichment analysis of protein sets confirmed our results.

This powerful approach allowed us to identify the molecular and metabolic bases of integrated trait variation, and therefore has a broad applicability domain.

## 1 INTRODUCTION

Phenotypic diversity within the living world results from billions of years of evolution. Most evolutionary pressures like mutation, random genetic drift, migration and recombination shape phenotypic diversity by directly changing the genetic composition of populations. The effects of selection are more difficult to predict because fitness is determined by phenotype, which results from a complex interaction between genotype and the environment (Fisher, 1930). An additional layer of complexity results from the fact that life-history traits (Stearns, 1992) are the observable results of unobservable processes that occur at the cellular level. During the last decades, there has been a growing interest for a better understanding in evolutionary biology of the so-called genotype-phenotype map (see *e.g*. Wagner and Zhang (2011)). In parallel, novel profiling technologies and accurate high-throughput phenotyping strategies have led to the genome-scale characterization of genomic sequences as well as to the quantification of transcriptomic, proteomic and metabolomic data at the individual level. Linking cellular processes to observable phenotypic traits is becoming a new discipline in Biology, known as integrative biology.

Unicellular organisms are the model species of choice for integrative biology because most of their observable traits are the direct product of cell metabolism, without needing to take into account the complexity of tissues and organs as in multicellular organisms. Schematically, cells sense the environment and transfer the information via signal transduction chains that interact with gene regulation networks. Gene regulatory networks modulate transcription, translation and post-translational modifications in response to environmental signals, resulting in variations in protein abundances. Differential abundances of enzymatic proteins affect the fluxes of matter and energy that are related to phenotypic traits, including life-history traits and fitness. Thus, in unicellular organisms, five integration levels are usually considered: genomic, transcriptomic, proteomic (including post-translational modifications), metabolic and cellular (or observable trait level). The last level is the most integrated, and it encompasses a variety of traits that are more or less related to fitness.

While technical progress has now allowed for genomic, transcriptomic, proteomic and trait levels to be readily measurable in a high number of individuals, metabolic fluxes are still difficult to measure. Although Metabolic Flux Analysis, based on Nuclear Magnetic Resonance (NMR) and differential usage of radioactive isotopes, is powerful (Antoniewicz, 2015), it remains low throughput and cannot be applied to a high number of individuals. Technical developments in mass spectrometry have boosted metabolomics (Nicholson and Lindon, 2008) by enabling the characterization of the metabolome, i.e. the complete set of metabolites in a cell, tissue, organ or organism. However, the technique still suffers from standardization difficulties and does not allow for high-throughput quantitative comparisons (Riekeberg and Powers, 2017).

Taking advantage of recent advancements in genome-scale functional annotation, constraint-based metabolic models provide a mathematical framework that allows us to predict internal cellular fluxes from *a priori* knowledge of thermodynamic constraints on individual enzymatic reactions, steady state hypotheses and the genome-scale stoichiometry matrix of all metabolic reactions. The idea is that a given set of environmental conditions will drive a cell to a steady state during which internal metabolites stay at a constant concentration while exchange fluxes are constant and correspond to a constant import/export rate. However, because the number of metabolites is much higher than the number of reactions, the system has an infinite number of solutions. Flux Balance Analysis (Fell and Small, 1986; Watson, 1984) consists in choosing, among all possible solutions, the solution that maximizes the biomass pseudo-flux that represents the conversion rate of biomass precursors into biomass. From a population genetics point of view, this method is questionable because evolution is not always based on optimization principles (Gould and Lewontin, 1979). However, it has been shown to be relevant in some cases, such as chemostat cultures of *Escherichia coli* (Edwards et al., 2001). Data-driven methods have also been proposed, which consist in choosing the most likely solution given observed transcriptomic, proteomic or metabolomic data (reviewed by Töpfer et al. (2015)). Among all available methods, the one from Lee et al. (2012) seems promising for studies at the population/species level. It is based on the realistic assumption that, at the genome scale, fluxes should covary with enzymatic protein abundances. Irrespective of the method, comparisons rely on the probability distribution of the solution space, which is analytically intractable because of the stoichiometry constraints. Recently, Braunstein et al. (2017) have proposed a Bayesian probabilistic method to characterize the solution space that is much faster than the classical Hit and Run algorithm (Bélisle et al., 1993) and allows for analyses at both the genome and population scale.

The HeterosYeast project investigated the molecular bases of heterosis in yeast at two different levels of integration: the proteomic level and the observable trait level (Blein-Nicolas et al., 2013, 2015; da Silva et al., 2015). A diallel study of two yeast species involved in wine fermentation was carried out and the hybrid and parental strains were monitored during fermentation of grape juice at two temperatures. Observable and proteomic traits were analyzed separately. In brief, the most important findings were: homeostasis of the interspecific hybrids observed at the trait level (da Silva et al. (2015)) and the predominance of interspecific heterosis at the proteomic level (Blein-Nicolas et al. (2015)). A closer analysis of genetic variance components confirmed that observable phenotypic traits tended to exhibit higher additive genetic variance and lower interaction variance than proteomic traits (Petrizzelli et al., 2019). However, the link between variation at the trait level and variation at the proteomic level is still missing.

Given the important genomic resources in yeast (Cherry et al., 2012a), a number of curated genome-scale metabolic models are now available (Caspi et al., 2014). Among these, the DynamoYeast model (Celton et al., 2012) describes central carbon metabolism in yeast. It is small enough (70 reactions and 60 metabolites) to remain tractable, and has been tested against experimental data (Nidelet et al., 2016).

The availability of the HeterosYeast dataset, combined with a curated metabolic model of central carbon metabolism in yeast and a probabilistic approach to explore the solution space, encouraged us to integrate experimental proteomic data into the metabolic model in order to predict unobserved metabolic fluxes. We used these predicted fluxes to bridge the gap between proteomic data and observable traits, and better understand the metabolic basis of life-history trait variation. This approach allowed us to show that the negative relationship between growth/fermentation traits and production traits is accounted for by a differential usage of the energy production pathway.

## 2 RESULTS

The HeterosYeast dataset provided valuable observations on the genetic diversity of yeast strains involved in the wine-making process at different levels of cellular organization, *i.e*. phenotypic traits related to life-history or fermentation (da Silva et al., 2015), and quantitative proteomic data (Blein-Nicolas et al., 2015). All traits were estimated or measured at 18°C and 26°C on a half-diallel design comprising 7 strains of *S. cerevisiae* and 4 strains of *S. uvarum*, with a total of 127 strain × temperature combinations.

In order to access an intermediate level of integration between protein abundances and traits, we used the DynamoYeast model, a curated Constraint-Based Model (CBM) of central carbon metabolism in yeast consisting of 70 reactions and 60 metabolites (Celton et al. (2012); Fig 1). Using gene-protein-reaction associations, enzymatic proteins and protein complexes were linked to the reactions of the CBM. Among the 70 enzymatic proteins and protein complexes, the abundances of 33 of them were retrieved from the dataset of 615 protein abundances quantified in the HeterosYeast project. Thus, the metabolic fluxes that best matched the observed patterns of variation of enzymatic protein abundance were retained (see Material and Methods).

**FIGURE 1.**
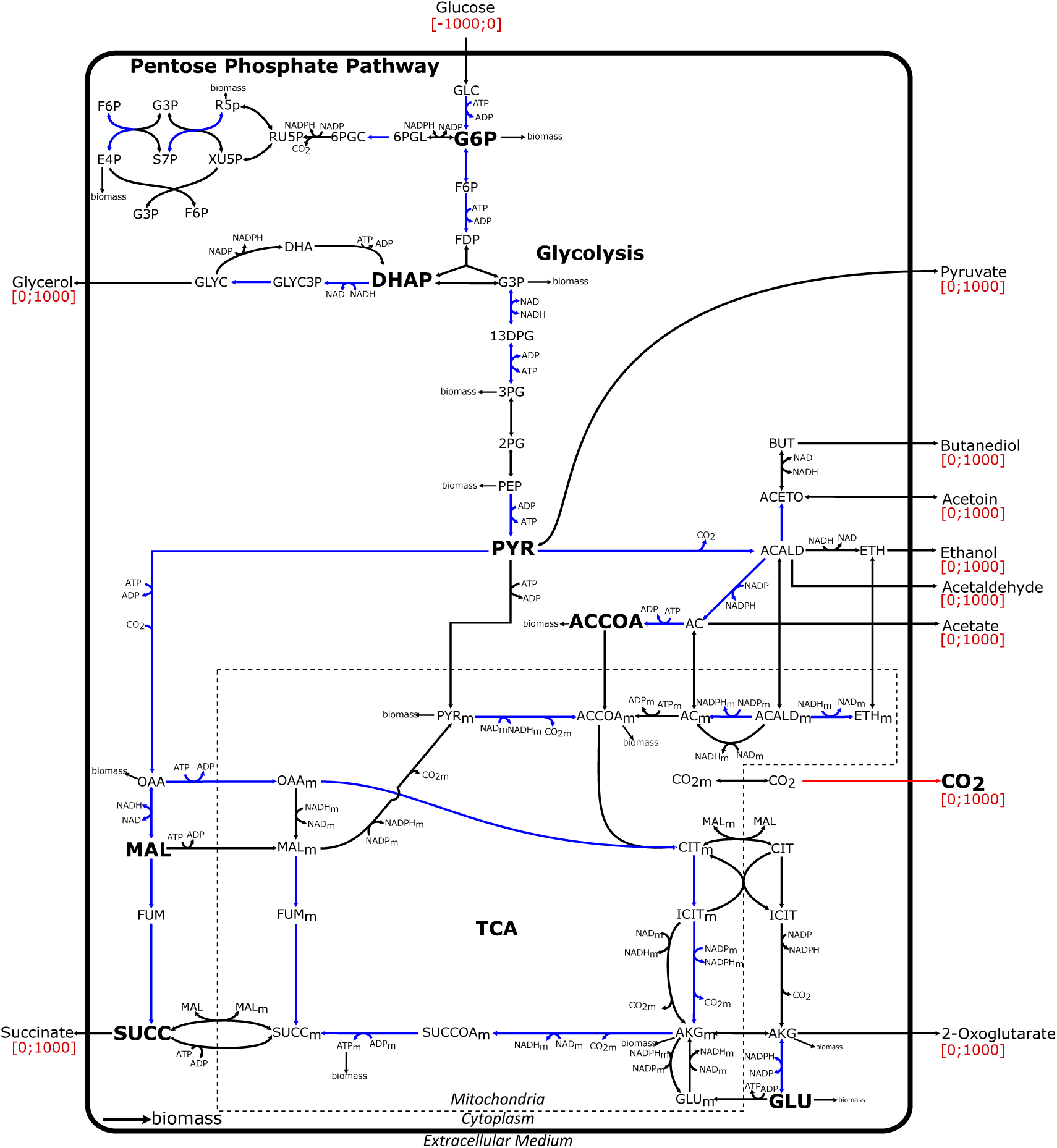
Representation of the DynamoYeast model of central carbon metabolism in *S. cerevisiae*. Metabolites are in black. Constraints on exchange fluxes are indicated in red between square brackets and correspond to fermentation, with glucose as the only input flux. The reaction color code is as follows: (*i*) in red, the experimentally measured CO_2_ exchange flux; (*ii*) in blue, the reactions associated with enzymatic proteins quantified in this study; (*iii*) in black, the other reactions present in the DynamoYeast model.

In brief, the strategy we proposed was: *(i)* to characterize the feasible solution space of the DynamoYeast model, *L*, through the posterior density distribution of the fluxes given by the Expectation Propagation algorithm (hereafter denoted EP, Braunstein et al. (2017)); *(ii)* to select a unique solution through minimization of the objective function that measures the Euclidean distance between observed enzyme abundances and reaction rates (*Z*, equation 11 in Material and Methods).

Below, we first describe the method and its validation using simulated datasets. Then, we analyze the relationship between the different integration levels, using the HeterosYeast dataset and the predicted fluxes of central carbon metabolism.

### 2.1 Sampling the feasible solution space with the Expectation Propagation algorithm

Sampling points from the feasible solution space *L* can be performed directly from the posterior truncated multivariate normal distribution of the fluxes. We compared the Hit and Run (HR) algorithm (Meersche et al., 2009) with the EP posterior distribution of the fluxes to test the goodness of prediction of the EP algorithm on the DynamoYeast posterior distribution. The EP methodology gave a good approximation of the mean and variance of the posterior marginal distribution of the fluxes (Supplementary Methods and Appendix Fig S1-S2), as well as of the variance-covariance matrix between fluxes (Appendix Fig S3). These results are similar to the ones obtained by Braunstein et al. (2017). Therefore, we decided to use the EP algorithm to sample the feasible solution space of the CBM.

### 2.2 Protein abundances are good predictors of the initial set of metabolic fluxes

Computer simulations were performed to assess the goodness of prediction of the proposed method, as detailed in section Testing the prediction algorithm. The two main parameters that were tested were: *(i)* the number of sampled points *N*_*s*_ in *L*; *(ii)* the number of observed proteins *N* ^obs^ to be included in the objective function *Z* (equation 11 in Material and Methods). To this end, a vector of flux values, ***v***^initial^, was first sampled from the feasible solution space of the DynamoYeast model. We then computed protein abundances by assuming different functional relationships between fluxes and enzymatic proteins, and we used the proposed method to predict metabolic flux values, ***v***^predicted^. Different *a priori* functional relationships between fluxes and enzymatic proteins led to similar results. Simulations showed that minimization of *Z* led to a high correlation between ***v***^initial^ and ***v***^predicted^ (Fig 2A). Correlations ranged from 0.65 to 0.99 (p-value < 0.05). By increasing the number of sampled points, *N*_*s*_ in *L*, the mean correlation slightly increased and its variance decreased. The number of observed protein abundances *N* ^obs^ had a more complex influence on the accuracy of the predictions. When increasing *N* ^obs^, the correlation between ***v***^initial^ and ***v***^predicted^ either increased, decreased or stayed constant, as illustrated in Fig 2B. However, the order of magnitude of the variation was small, and the correlation tended to be more stable for high *N*_*s*_ value (Fig 2B). When considering the actual number of enzyme abundances (*N* ^obs^ = 33) that were matched between the HeterosYeast proteomic data and the DynamoYeast CBM, we observed a high correlation between ***v***^initial^ and ***v***^predicted^ after setting *N*_*s*_ = 10^6^ (Fig 2C). Altogether, we considered that our algorithm was efficient for predicting unobserved fluxes from enzyme abundances, given the structure of the metabolic network.

**FIGURE 2.**
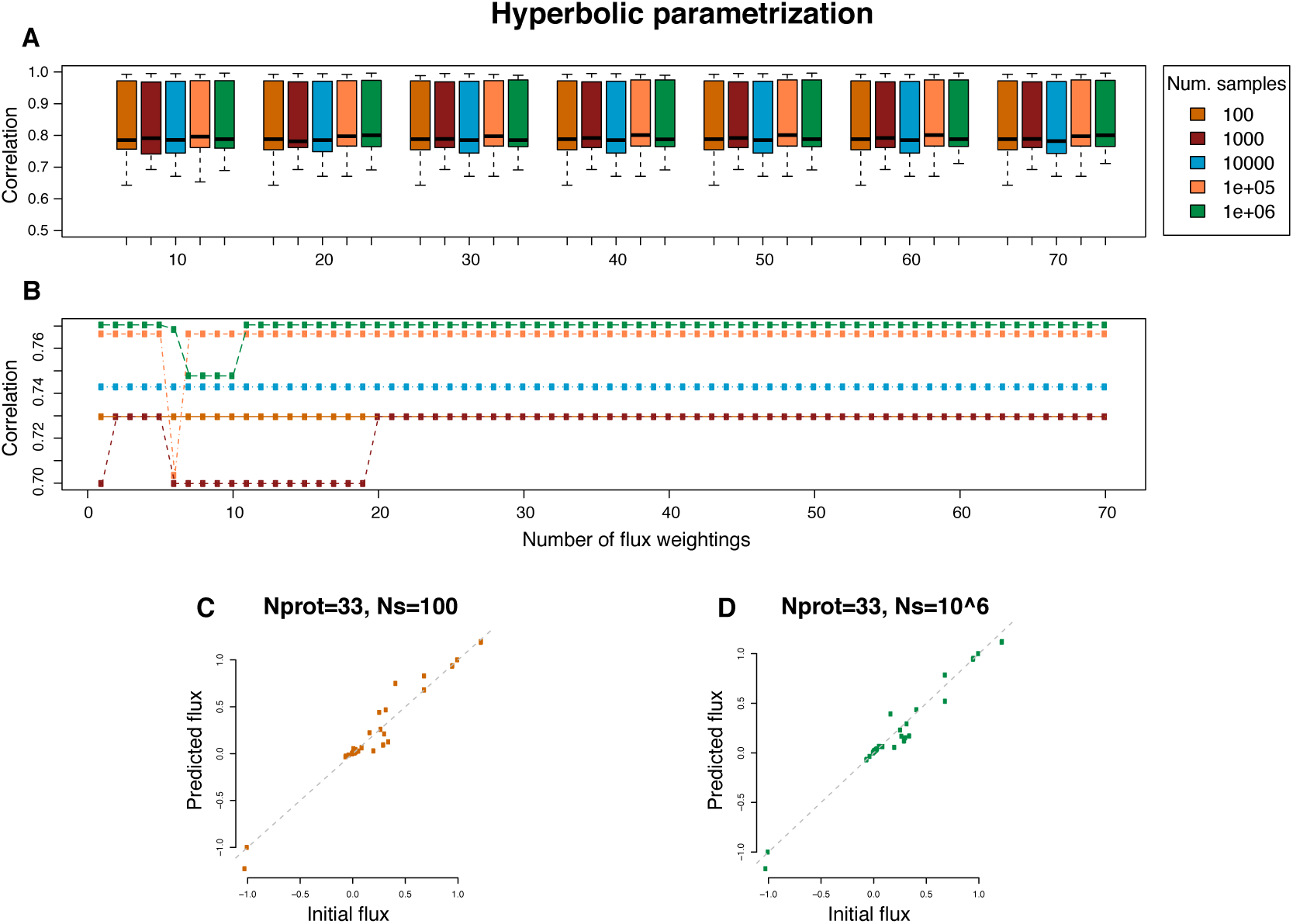
Correlation between initial and predicted fluxes in simulated datasets using the DynamoYeast model. Enzymatic protein abundances were expressed in terms of a hyperbolic function of the initial fluxes using equation 12. Colors indicate the number of points *N*_*s*_ that were sampled in solution space *L*. **A**. Boxplot representation as a function of the number of observed proteins *N* ^obs^, from 10 to 70, in increments of ten. Each box represents 1,000 simulations. **B**. Observed changes in the correlation during a single simulation run when the number of observed proteins was increased in increments of 1, from 1 to 70 (same color code as in **A**). **C**. The relationship between initial and predicted fluxes shown for one simulation with *N* ^obs^ = 33 and *N*_*s*_ = 100. **D**. The relationship between initial and predicted fluxes shown for one simulation *N* ^obs^ = 33 and *N*_*s*_ = 10^6^.

### 2.3 Predicting unobserved fluxes from the observed variation in protein abundances

The HeterosYeast proteomic data were used in the context of the DynamoYeast model of central carbon metabolism in yeast. In addition, for each strain × temperature combination, the observed CO_2_ release rate was used as an additional constraint in the form of *a priori* knowledge to delimit the feasible solution space *L*. We sampled *N*_*s*_ = 10^6^ points in *L* to select a unique solution that minimizes the Euclidean distance between fluxes and enzyme abundances. As a result, we predicted 69 unobserved fluxes in the CBM for each of the 127 strain × temperature combinations. Statistical approaches were then used to investigate the variation components and the structure of the new dataset *D* consisting of 615 protein abundances (**E**), 70 metabolic fluxes (**V**) and 28 fermentation and life-history traits (**T**):

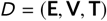

### 2.4 Patterns of variation depend on the integration level

The 127 observations in the new dataset *D* had a specific structure. There were 7 parental strains (*S.c*.) and 21 intraspecific hybrids from *S. cerevisiae* (*S.c*.×*S.c*.), 4 parental strains (*S.u*.) and 6 intraspecific hybrids from *S. uvarum* (*S.u*.×*S.u*.), and 28 interspecific hybrids (*S.c*.×*S.u*.). All strains were observed during alcoholic fermentation of wine grape juice at two temperatures, 18°C and 26°C (da Silva et al., 2015).

To better understand the patterns of variation at each integration level, Principal Component Analyses (PCA) were performed for each type of trait separately. Results are presented in Fig 3, where strains are identified by species, type of cross (intraspecific hybrid, interspecific hybrid or parental strain) and temperature. The first PCA components accounted for 20%, 23% and 27% of the total variation and the second for 14%, 18% and 19% of the total variation for protein abundances, metabolic fluxes and fermentation/life-history traits, respectively (Fig 3A, B and C). Different integration levels displayed different patterns of phenotypic diversity.

**FIGURE 3.**
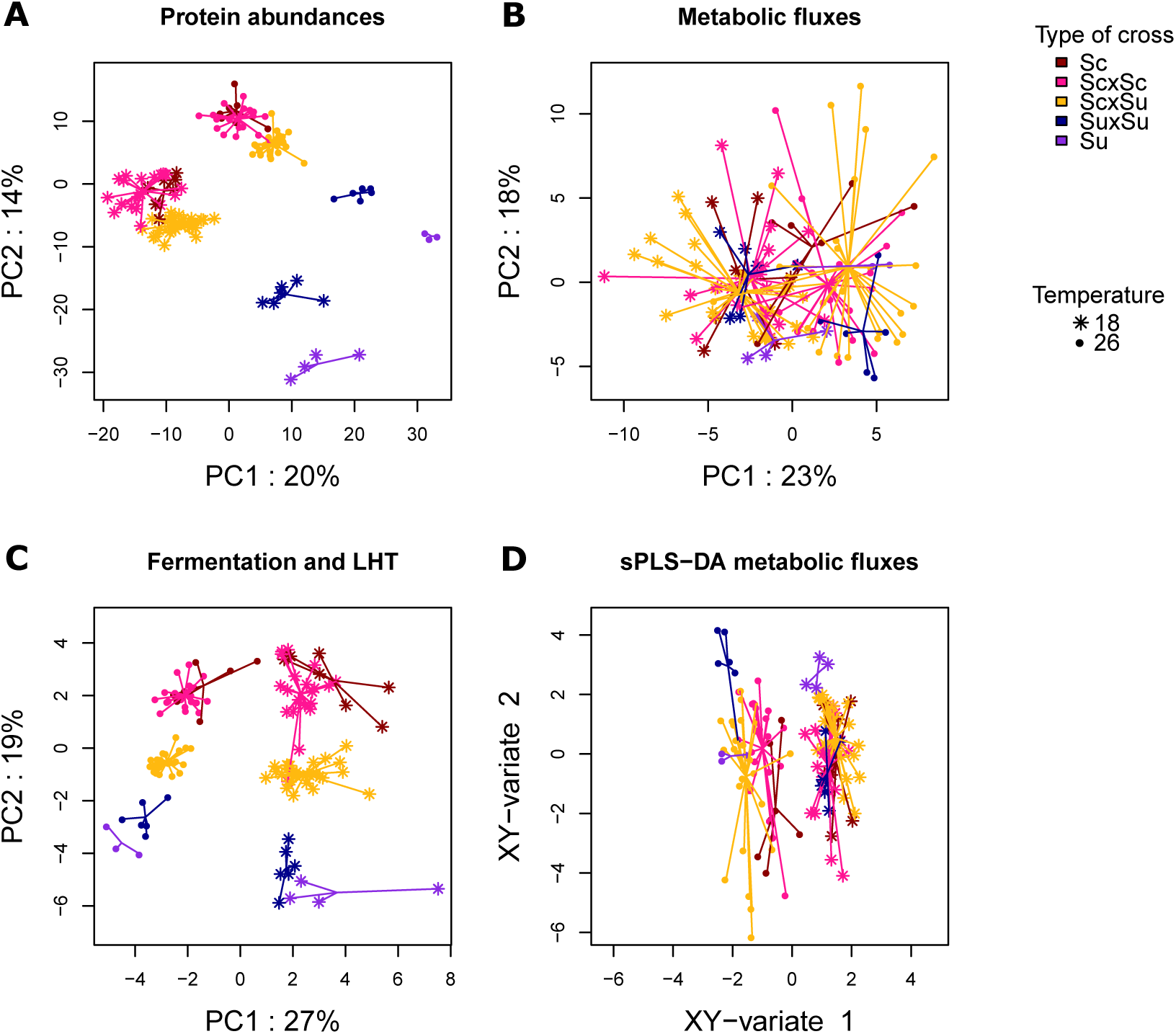
Principal Component Analysis and sparce Partial Least Square-Discriminant Analysis. PCA for protein abundances (A), metabolic fluxes (B) and fermentation/life-history traits (C). sPLS-DA for metabolic fluxes (D). Observations are represented on the first two PCA axes (sPLS-DA, respectively). Each dot corresponds to a strain × temperature combination. Different types of dots correspond to different temperatures, while the type of cross is color-coded. Segments join the data points to the centroid of the group (cross × temperature) to which they belong.

At the proteomic level (**E**), the first two PCA axes contributed to both differences between temperatures and between species and type of cross. Heterosis was observed for the three types of hybrids at both temperatures. First, *S.u*.×*S.u*. hybrids were clearly differentiated from their *S.u*. parents. Second, *S.c*.×*S.u*. interspecific hybrids were closer to their *S.c*. parents than their *S.u*. parents. Finally, *S.c*.×*S.c*. hybrids were close to their *S.c*. parents, but the range of variation between *S.c*.×*S.c*. hybrids was greater than between parental strains. Altogether, protein abundance in a hybrid strain cannot be predicted by the mean of its parental values.

At the trait level (**T**), we observed a high temperature effect, with axis 1 (27% of the variation) clearly separating the strains that were grown at 26°C from those that were grown at 18°C. At 26°C, strains were characterized by high growth rates (*r*), high CO_2_ fluxes (*J*_max_ and *V*_max_), high *Hexanol* and *Decanoic acid*, a low carrying capacity (*K*) and short fermentation times (*AFtime, t-lag, t-75, t-45*, Fig S4). At 18°C, strains were characterized by low growth rates and low CO_2_ fluxes, a high *K* and long fermentation times (Fig S4). These two groups of traits mostly varied with temperature, although some differences between strains were observed within rather than between types of cross, especially at 18°C. At 26°C, *S.u*. strains perform slightly better than *S.c*. strains (higher growth rates, faster fermentation). The types of cross were clearly separated along PCA axis 2. Heterosis was observed at the trait level in intraspecific hybrids. However, interspecific hybrids seemed to be in-between the two parental strains. Traits that explain the differences between the observations along axis 2 were cell size (*Size-t-N*_max_) and *Ethanol* at the end of fermentation (positively correlated with axis 2), aroma production at the end of fermentation, as well as *Sugar.Ethanol.Yield* (negatively correlated) (Fig S4). Note that these traits were not influenced by temperature. Hence, at the trait level, we observed differences between yeast species for traits related to aroma production that were not influenced by temperature. Most fermentation and life-history traits showed a strong temperature effect, large differences between strains within a type of cross and weak heterosis.

At the flux level (**V**), temperature separated the observations on axis 1, but both axis 1 and axis 2 distinguished strains independently of their origin. Notice however that the variation range in hybrids was greater than in the parental strains, indicating differences between inbred and hybrid strains. Altogether, central carbon metabolic fluxes were influenced by temperature and showed strong differences between strains that were not related to the type of cross or the parental species. Sparse Partial Least Squares Discriminant Analysis (sPLS-DA) was performed on metabolic fluxes in order to select the main features that characterize the species × temperature combinations (Fig 3D). As previously, the first axis distinguished strains grown at different temperatures. Six fluxes contributed to the first axis of the sPLS-DA: CO_2_, ethanol, pyruvate decarboxylase, alcohol dehydrogenase, 6-phosphogluconolactonase and phosphogluconate dehydrogenase fluxes (Fig S5). All were negatively correlated with axis 1 and were involved in fermentation. This shows that fermentation was more efficient at 26°C. The second axis distinguished inbred strains from intraspecific hybrids with a genotype × temperature interaction: both *S.u*.×*S.u*. and *S.c*.×*S.c*. hybrids had higher coordinates than their parents at 26°C, whereas *S.u*.× *S.u*. had lower coordinates than their parents at 18°C, and *S.c*.×*S.c*. hybrids were confounded with their parental strains. Interspecific hybrids were characterized by a wide variation range at both temperatures. Fluxes that contributed to axis 2 were mainly mitochondrial fluxes. Mitochondrial acetyl-CoA formation, mitochondrial citrate synthase, mitochondrial aconitate hydratase, mitochondrial isocitrate dehydrogenase (NAD+) and mitocondrial transport fluxes of pyruvate, oxaloacetate and acetaldehyde were negatively correlated with the second axis, while mitochondrial transport of 2-oxodicarboylate, ethanol and CO_2_ fluxes were positively correlated (Fig S5).

In summary, we found at each integration level a strong temperature effect, large differences between strains, and evidence for heterosis, *i.e*. differences between hybrids and mid-parent values. However, patterns differed between the proteomic and the most integrated level. At the proteomic level, proteins involved in differences between strains were the same as the ones involved in differences between species and between temperatures. At the flux level, there were few differences between species. Differences between temperatures were associated with enzymatic reactions related to fermentation, while differences between strains were associated with enzymatic reactions that were either involved in fermentation, or in the part of the TCA cycle that takes place in mitochondria. At the trait level, differences between temperatures were associated with differences in growth and fermentation traits, which were relatively conserved within species but showed between-strain variation. Differences between species mostly concerned volatile compounds that are produced by secondary metabolism at the end of fermentation.

### 2.5 Fermentation and life-history traits are associated with different metabolic pathways of carbon metabolism in yeast

Regularized Canonical Correlation Analysis (rCCA) was performed to investigate the correlation between metabolic fluxes and fermentation/life-history traits (Fig 4). Fermentation and life-history traits could be divided into two main groups showing contrasting profiles. The first group consisted of traits that clustered with the carrying capacity, *K*. They were negatively correlated with fluxes involved in the glycolysis, ethanol synthesis and pentose phosphate pathways, and positively correlated with fluxes in the TCA reductive branch. By contrast, the second group consisted of traits that clustered with the intrinsic growth rate, *r*, and were positively correlated with fluxes involved in the glycolysis, ethanol synthesis and pentose phosphate pathways and negatively correlated with fluxes in the TCA reductive branch. Consistent with these results, the *biomass* pseudo-flux was positively correlated with *r* and negatively correlated with *K*.

**FIGURE 4.**
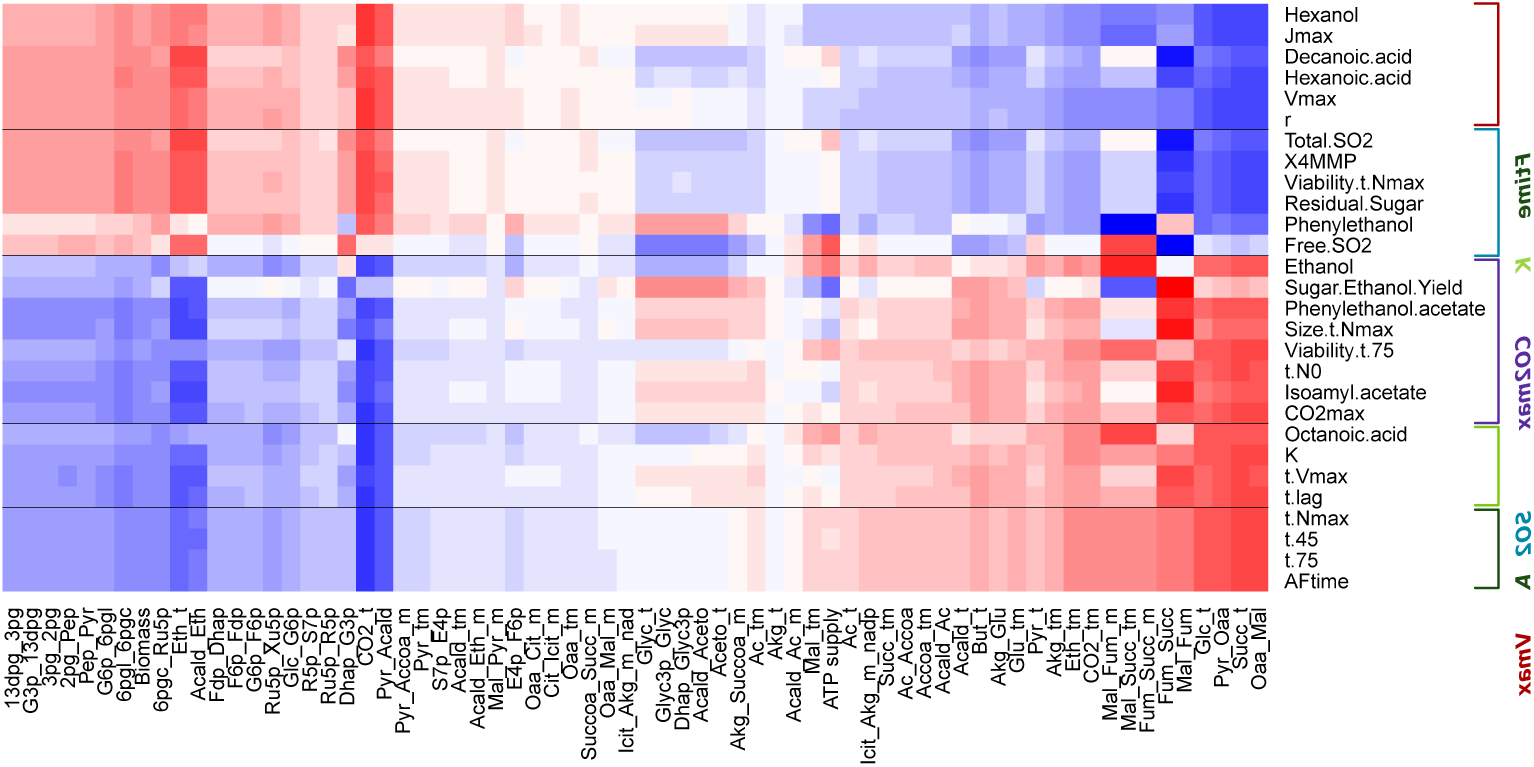
Regularized Canonical Correlation Analysis of metabolic fluxes and fermentation/life-history traits. Penalization parameters were tuned through a leave-one-out cross-validation method on a 1000 × 1000 grid between 0.0001 and 1 (*λ*_1_ = 0.8, *λ*_2_ = 0.0001). Canonical correlation values between metabolic fluxes and fermentation/life-history traits are represented as a gradient of colors from blue (−1) to red (+1). Metabolic fluxes (columns) and fermentation/life-history traits (rows) were clustered using the *hcl ust* method. Flux names are encoded as abbreviations of the substrate and the product of the reaction connected by “_”. The five groups defined by fermentation and life-history traits are shown on the right.

When looking at the flux correlation structure described in Fig 4, we see the opposition between glycolytic fluxes and TCA fluxes. High growth rates and CO_2_ fluxes (*J*_max_, *V*_max_) and correspondingly fast fermentation (short fermentation times) seem to be associated with a central carbon metabolism oriented towards fermentation, while high carrying capacity, low growth rate and slow fermentation seem to be associated with a central carbon metabolism oriented towards the production of secondary metabolites (succinate, pyruvate, acetate, acetaldehyde and butanediol).

The *K* group could be divided into three subgroups, based mainly on the correlation between traits and the glycerol synthesis and acetaldehyde fluxes: **AFtime, K** and **CO**_**2**_**max** (subgroups designated by the name of the main trait in boldface). The **AFtime** subgroup showed a slightly negative correlation, the **K** subgroup a slightly positive correlation and the **CO**_**2**_**max** subgroup a positive correlation. **AFtime** grouped most traits correlated with the duration of fermentation, *AFTime, t-45, t-75, t-N*_max_; **K** grouped traits measuring the lag time and the beginning of fermentation (*t-lag, t-V*_max_), the carrying capacity (*K*) and the concentration of *Octanoic acid* (a fatty acid) at the end of fermentation, while **CO**_**2**_**max** grouped traits correlated with fermentation products (*total* CO_2_, *Ethanol* and *Sugar.Ethanol.Yield*), two volatile esters, *Isoamyl acetate* and *Phenyl-2-ethanol acetate*, as well as cell size and cell viability measured close to the end of fermentation, and *t-N*_0_.

Similarly, within the *r* group we distinguished two clusters of traits: **SO**_**2**_ and **Vmax. SO**_**2**_ grouped basic enological parameters measured at the end of fermentation (*total* and *free SO*_2_, *residual sugar*), cell viability measured once carrying capacity is reached (*Viability-t-N*_max_), and two volatile compounds *Phenyl-2-ethanol* (alcohol) and *4-methyl-4-mercaptopentan-2-one* (*X4MMP*, thiol). **Vmax** grouped traits that correlated with *V*_max_ and *r*, such as *J*_max_ and the amount of *hexanol* (alcohol) and *hexanoic* and *decanoic acids* (fatty-acid) that were quantified at the end of fermentation (Table 1).

**TABLE 1.**
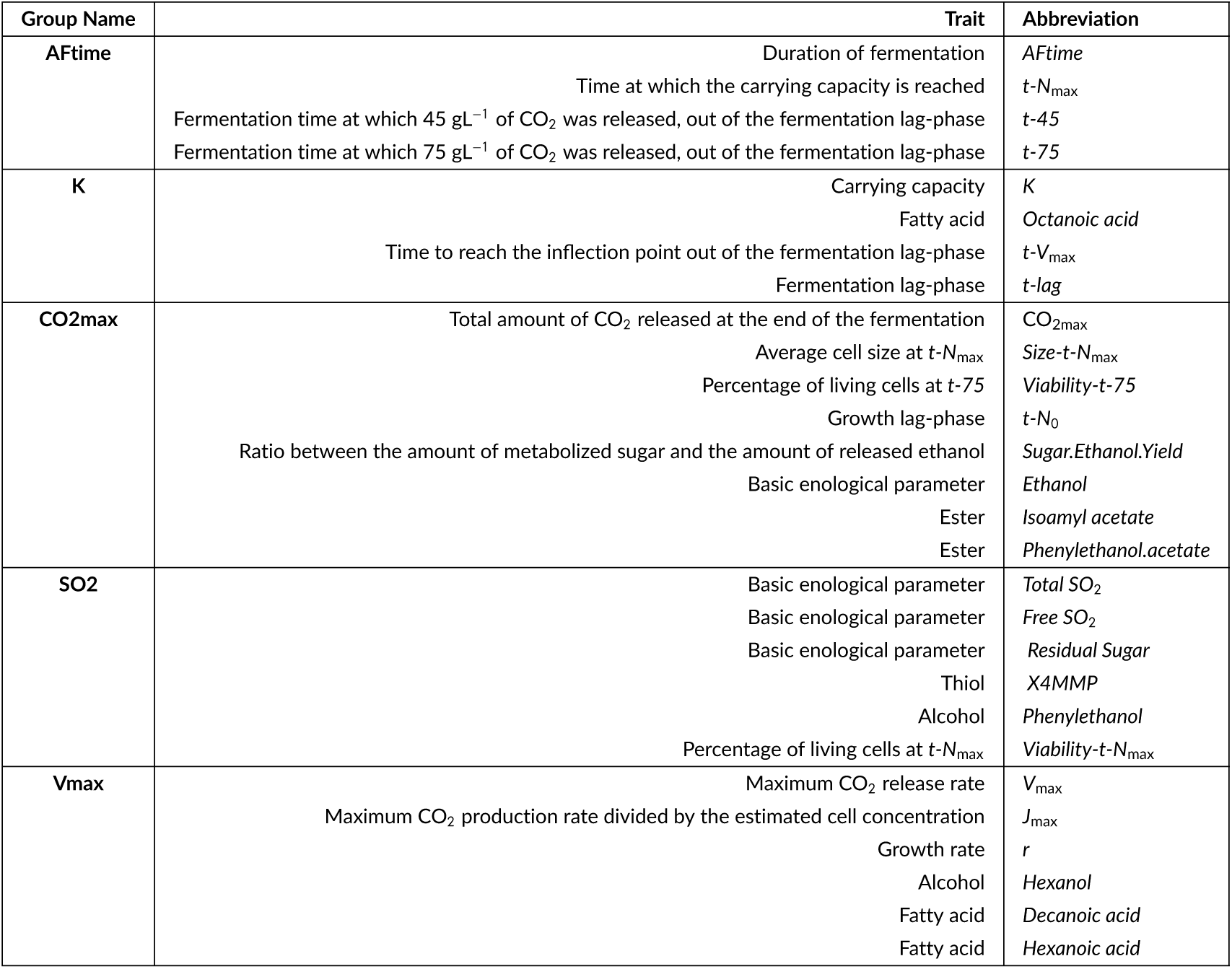
Group name, trait description and abbreviation for each fermentation/life-history trait analyzed in this study. Traits are grouped following the correlation structure obtained by rCCA (Fig 4).

| Group Name | Trait | Abbreviation |
| --- | --- | --- |
| **AfTime** | Duration of fermentation | *AfTime* |
| | Time at which the carrying capacity is reached | $t_{-N_{\max}}$ |
| | Fermentation time at which 45 gL<sup>-1</sup> of CO<sub>2</sub> was released, out of the fermentation lag-phase | $t_{-45}$ |
| | Fermentation time at which 75 gL<sup>-1</sup> of CO<sub>2</sub> was released, out of the fermentation lag-phase | $t_{-75}$ |
| **K** | Carrying capacity | $K$ |
|  | Fatty acid | *Octanoic acid* |
| | Time to reach the inflection point out of the fermentation lag-phase | $t_{-V_{\max}}$ |
| | Fermentation lag-phase | $t_{\text{-lag}}$ |
| **CO<sub>2</sub>max** | Total amount of CO<sub>2</sub> released at the end of the fermentation | $\text{CO}_{2\max}$ |
| | Average cell size at $t_{-N_{\max}}$ | $\text{Size-}t_{-N_{\max}}$ |
| | Percentage of living cells at $t_{-75}$ | $\text{Viability-}t_{-75}$ |
| | Growth lag-phase | $t_{-N_0}$ |
|  | Ratio between the amount of metabolized sugar and the amount of released ethanol | *Sugar.Ethanol.Yield* |
|  | Basic enological parameter | *Ethanol* |
|  | Ester | *Isoamyl acetate* |
|  | Ester | *Phenylethanol.acetate* |
| **SO<sub>2</sub>** | Basic enological parameter | *Total SO<sub>2</sub>* |
|  | Basic enological parameter | *Free SO<sub>2</sub>* |
|  | Basic enological parameter | *Residual Sugar* |
|  | Thiol | *X4MMP* |
|  | Alcohol | *Phenylethanol* |
| | Percentage of living cells at $t_{-N_{\max}}$ | $\text{Viability-}t_{-N_{\max}}$ |
| **Vmax** | Maximum CO<sub>2</sub> release rate | $V_{\max}$ |
| | Maximum CO<sub>2</sub> production rate divided by the estimated cell concentration | $J_{\max}$ |
| | Growth rate | $r$ |
|  | Alcohol | *Hexanol* |
|  | Fatty acid | *Decanoic acid* |
|  | Fatty acid | *Hexanoic acid* |

In summary, we were able to associate fermentation and life-history traits with metabolic fluxes based on their correlation patterns. In particular, we found that the negative correlation between *r* and *K* is explained by a different pathway usage of the central carbon metabolism. A high *r* and a low *K* are associated with glycolysis and fermentation, while a low *r* and a high *K* are associated with the TCA cycle and respiration.

### 2.6 Metabolic bases of phenotypic trait variation in yeast

In order to confirm the association between integrated trait variation and the differential usage of central carbon metabolism, we identified proteins involved in trait patterning that were not included in the DynamoYeast model, as observed from the correlations between traits and fluxes (see section Statistical Analysis). We performed a Linear Discriminant Analysis on the correlation matrix between the **T** traits and the **E** proteins using as discriminant features the five groups of fermentation and life-history traits described above.

Linear Discriminant Analysis clearly separated the five trait categories along the first axis, which explains 99% of the total variation (Fig 5). **AFtime** and **K** traits were close to each other, and had positive coordinates on LDA1; **Vmax** had high negative coordinates, **SO**_**2**_ had a slightly negative mean and **CO2max** had a slightly positive mean on LDA1. Given the high discriminative power of LDA1, it is clear that proteins that were positively or negatively correlated with LDA1 participate in the differentiation of **AFtime** and **Vmax** trait groups.

**FIGURE 5.**
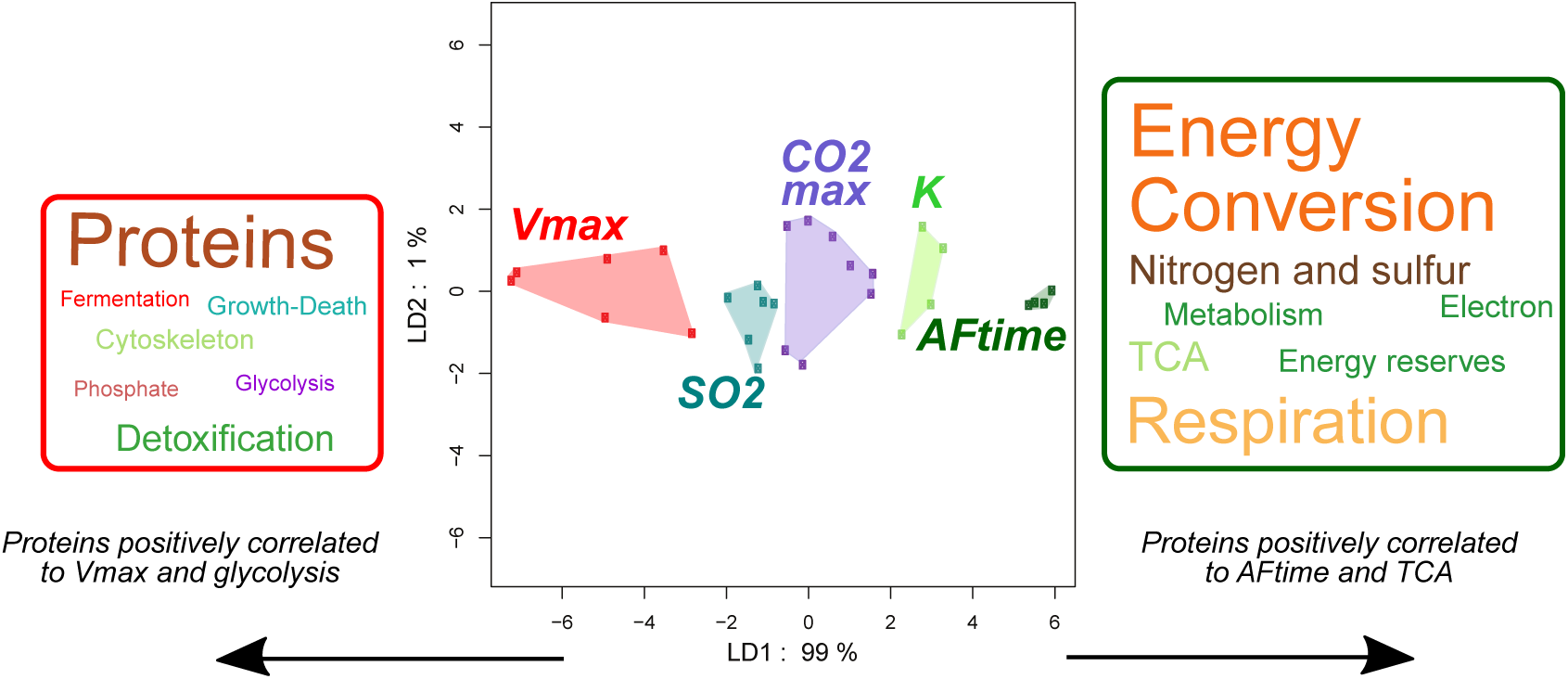
Projection of the 28 fermentation/life-history traits onto the first two axes of a Linear Discriminant Analysis of protein abundances. Groups of traits were defined from their correlation with central carbon metabolism fluxes. Each dot corresponds to one fermentation or life-history trait. Colors correspond to groups of traits, which are named after one representative trait. The results confirm the structure of fermentation and life-history traits and reveal two groups of traits with antagonistic proteomic patterns: the **AFtime** group and the **Vmax** group. The functional enrichment of proteins that are positively or negatively correlated with the first axis is represented by a cloud of words.

Functional analysis of proteins that best correlate with the first axis of the LDA was performed on the group of proteins with a correlation of 0.85 in the positive and in the negative direction. Pearson’s chi square test of enrichment showed that the group of proteins that were negatively correlated with the first axis was enriched in proteins linked to protein fate, cytoskeleton, detoxification, growth and death but also to the fermentation, glycolysis and phosphate pathways. The group of proteins that were positively correlated with LDA1 was enriched in proteins linked to energy conversion, nitrogen and sulfur pathways, metabolism, energy reserves, electron transport and respiration. This result was represented as a cloud of words in Fig 5.

In conclusion, the association between trait variation and central carbon metabolism observed at the flux level is confirmed by the proteomic analysis. Proteins that covary with traits of the **Vmax** group and with glycolytic and fermentation fluxes are involved not only in glycolysis and fermentation, but also in protein synthesis and degradation (protein fate) and in cytoskeleton formation, which can be associated with cell division. Proteins that covary with traits of the **AFtime** group and with the TCA cycle and respiration fluxes are involved not only in the TCA cycle and respiration, but also in electron transport, energy conversion and nitrogen and sulfur metabolism.

## 3 DISCUSSION

We applied cutting-edge methods of data integration to an original yeast dataset. The HeterosYeast dataset comprised quantitative proteomic data as well as fermentation and life-history traits measured during wine fermentation on a range of strains from two yeast species. The objective was to integrate information at different levels of cellular organization (proteomic and metabolic fluxes) to better understand the metabolic bases of phenotypic variation in yeast, in particular life-history traits related to fitness. The key point of this study was to incorporate proteomic data in a constraint-based metabolic model to estimate the values of unobserved metabolic fluxes. Using a combination of multivariate analyses of heterogeneous high-dimensional datasets, we were able to show that the metabolic flux level retains information that is not directly interpretable at the proteomic or trait level. In particular, we showed that the negative correlation between traits associated with population growth rate and traits associated with maximum population size (carrying capacity) could be explained by a differential usage of central carbon metabolism, in this case fermentation *versus* TCA cycle.

### 3.1 Constraint-based modeling can predict unobserved fluxes from observations at the cellular level

Functional genome annotations, coupled with current knowledge in biochemistry, now allow cell metabolism to be described at the genome scale, using constraint-based metabolic models that take into account the stoichiometry of each reaction and incorporate thermodynamic constraints (Palsson, 2015). Without any *a priori* knowledge, the number of steady-state solutions for reaction rates is infinite; however, this number can be reduced by integrating observations. Three types of experimental data can be used: *(i)* exchange metabolic fluxes; *(ii)* metabolite input/output rates and *(iii)* protein abundances. External metabolic fluxes and metabolite input/output rates can be used directly in constraint-based models to reduce the feasible solution space, *L* (equation 5 and inequality 6) under the steady-state assumption.

Protein abundances, linked to the metabolic fluxes considered in the model through a gene-protein-reaction (GPR) association, carry information regarding network functioning and the state of the metabolic network at a given time and under specific conditions. Following Lee et al. (2012), we used protein abundance profiles to find the set of metabolic fluxes that minimized the Euclidean distance between metabolic fluxes and enzyme abundances. Indeed, even though the relationship between flux and enzyme abundances is commonly non-linear, the extent to which a particular pathway is used is more or less associated with the abundance of its enzymes (Sabarly et al., 2016).

The method described here relies on a probabilistic approach. Following Braunstein et al. (2017), we chose to characterize the feasible solution space *L* by means of its posterior density distribution calculated with the Expectation Propagation (EP) algorithm. The computation time of the EP algorithm is much shorter than that of the well-known Hit and Run algorithm (Bélisle et al., 1993), allows to sample metabolic fluxes in *L* and provides their associated posterior probability. In order to select a unique solution in *L*, we minimized *Z*, the Euclidean distance between observed protein abundances and the associated metabolic fluxes weighted by the inverse of the probability of observing such a set of fluxes, *p*_***v***_ (equation 11). This minimization process involved sampling in *L*, and selection was made after computing *Z* over a high number of sampled points.

Computer simulations confirmed that our method had good prediction efficiency. In particular, we showed that prediction efficiency was not affected by the non-linearity of the flux-enzyme relationship. The most important parameter was the number of reactions *N* ^obs^ for which proteomic observations were available, compared to the CBM size, *n*. When *N* ^obs^ was too low, adding new information led to a decrease in prediction efficiency. A decrease in correlation between initial and predicted fluxes means that, when a new enzyme is added, the solution that minimized the total Euclidean distance can lead to flux predictions that are farther from their true value. This can occur whenever there is a weak correlation between the first *n* − 1 fluxes, and the additional flux *v*_*n*_. Therefore, it is important that protein abundance observations cover the main features of the architecture of the metabolic network. Here, we observed protein abundances for 33 out of the 70 reactions of the DynamoYeast model, which was sufficient to attain high prediction accuracy. Recent progress in gel-free/label-free quantitative proteomics now allows us to quantify thousands of proteins and should ensure good coverage even for metabolic models at the genome scale (Belouah et al., 2019).

Even though our flux predictions would need to be verified, we are confident that our method uncovers the main processes of cell metabolism. Indeed, our method integrates additional information about the known architecture of the metabolic network to predict unobserved fluxes from observed protein abundances and globally adds information to the system.

### 3.2 Unraveling the metabolic bases of life-history trait variation

In this study, we predicted the metabolic fluxes of central carbon metabolism in a population obtained from a half-diallel cross between two species of yeast, *S. cerevisiae* and *S. uvarum*. The genetic values of 615 protein abundances and 28 fermentation/life-history traits were estimated under fermentation conditions at two different temperatures, 18°C and 26°C, leading to a total of 127 observations from 66 different yeast strains (Albertin et al., 2013b). As described above, we predicted metabolic fluxes for each strain × temperature combination by coupling the DynamoYeast model, a highly curated Constraint-Based Model of central carbon metabolism (Celton et al., 2012) using the observed CO_2_ release rate as *a priori* knowledge, and measurements of protein abundances associated with 33 out of the 70 reactions in the model.

The final dataset consisted of three matrices of 127 × 615 protein abundances, 127 × 70 central carbon fluxes, and 127 × 28 fermentation/life-history traits. As the total number of characters (713) greatly exceeded the number of samples (127), we used regularization techniques for the multivariate analyses (Rohart et al., 2017). In order to relate the variation patterns we observed at different integration levels, we used a top-down strategy from the most integrated to the least integrated level. First, we explored the correlation between traits and metabolic fluxes. Second, we identified the proteins which were not included in the metabolic model that best explained the correlation between traits and fluxes.

In our dataset, we found a negative correlation between traits associated with growth and CO_2_ fluxes, and traits associated with population size and fermentation length. These negative correlations reflected different life-history strategies, as has been observed previously in different yeast collections from either industrial (Albertin et al., 2013a) or natural sources (Spor et al., 2008, 2009). This broadly corresponds to the well-known *r-K* trade-off in ecology (Pianka, 1970). More recently, Collot et al. (2018) suggested that such a trade-off could arise from eco-evolutionary feedback loops because competing strains also modify their environment through the production of different sets of metabolites. The HeterosYeast dataset shows that the choice of a strategy is plastic (da Silva et al., 2015) and can be modified by the environment (here the fermentation temperature).

By adding information to the DynamoYeast model, we showed that such a trade-off can be explained by metabolic switches between fermentation associated with glycolysis, and secondary metabolite production, associated with the TCA cycle. This duality in the functioning of yeast central carbon metabolism was observed by (Nidelet et al., 2016), who matched the DynamoYeast model to experimentally measured exchange fluxes in different *S. cerevisiae* strains. The switch between the two modes of functioning (Fig 4) depends partly on the balance between two isoforms of alcohol dehydrogenase (ADH). Interestingly, Albertin et al. (2013a) previously found that the trade-off between cell size and *K* is related to changes in the percentage of acetylation of ADH 1p, with high levels being associated with large cells and low *K*

Because this paper describes a proof of concept, we deliberately chose to focus on central carbon metabolism and we used the DynamoYeast model because it contains a small number of reactions compared to available genome-scale models (Caspi et al., 2014). Therefore, we were not able to explain between-strain variation for traits related to secondary metabolism like aroma production, which merely discriminated between the two yeast species. Moreover, only a small subset of the proteomic data was coupled with the metabolic model. By searching for the proteins that best explained the trait patterns revealed at the flux level, we were able to identify proteins that were associated with the *r-K* trade-off at the trait level. Analysis of protein functional annotations confirmed the known link between the glycolysis and pentose-phosphate pathways and fermentation, and between extensive usage of TCA cycle and energy conversion.

Altogether, by coupling phenomic data with mathematical modeling of metabolism and cutting-edge statistical analyses (taking into account the high-dimensionality and heterogeneity of the measures), we were able to explain a well-known trade-off between two sets of yeast life-history traits by the differential usage of energy production pathways. Glycolysis and fermentation lead to fast growth and resource consumption. TCA and secondary metabolite production lead to slow growth and high population size. The duality between the two alternative uses of the central carbon metabolism is encoded into the architecture of the metabolic network.

## 4 MATERIAL AND METHODS

### 4.1 Materials

#### 4.1.1 Materials: The HeterosYeast dataset

The genetic material for the experimental study consisted of 7 strains of *S. cerevisiae* and 4 strains of *S. uvarum* associated with various food processes (enology, brewing, cider fermentation and distilling) or isolated from the natural environment (oak exudates). The 11 parental lines were selfed and pairwise crossed, which resulted in a half-diallel design with a total of 66 strains: 11 inbred lines, 27 intraspecific hybrids (21 for *S. cerevisiae*, noted *S. c*. × *S. c*., and 6 for *S. uvarum*, noted *S. u*.× *S. u*.) and 28 inter-specific (noted *S. c*. × *S. u*). The 66 strains were grown in triplicate in fermentors at two temperatures, 26°C and 18°, in a medium similar to enological conditions (Sauvignon blanc grape juice, da Silva et al. (2015)). From a total of 396 alcoholic fermentations (66 strains, 2 temperatures, 3 replicates), 31 failed due to the poor fermenting ability of certain strains. The design was set up as a block of two sets of 27 fermentations (26 plus a control without yeast to check for contamination), one carried out at 26°C and the other at 18°. The distribution of the strains in the block design was randomized to minimize residual variance of the estimators of the strain and temperature effects, as described in Albertin et al. (2013a).

For each alcoholic fermentation, two types of phenotypic traits were measured or estimated from sophisticated data adjustment models: 35 fermentation/life-history traits and 615 protein abundances. Fermentation/life history traits were classified into four categories (da Silva et al., 2015):

- *Kinetics parameters*, computed from the CO_2_ release curve modeled as a Weibull function fitted on CO_2_ release quantification monitored by weight loss of bioreactors: the fermentation lag-phase, *t-lag* (h); the time to reach the inflection point out of the fermentation lag-phase, *t-V*_max_ (h); the fermentation time at which 45 gL^−1^ and 75 gL^−1^ of CO_2_ was released, out of the fermentation lag-phase, *t-45* (h) and *t-75* (h) respectively; the time between *t-lag* and the time at which the CO_2_ emission rate became less than, or equal to, 0.05 gL^−1^h^−1^, *AFtime* (h); the maximum CO_2_ release rate, *V*_max_ (gL^−1^*h*^−1^); and the total amount of CO_2_ released at the end of the fermentation, CO_2max_ (gL^−1^).
- *Life history traits*, estimated and computed from the cell concentration curves over time, modeled from population growth, cell size and viability quantified by flow cytometry analysis: the growth lag-phase, *t-N*_0_(*h*); the carrying capacity, *K* (log[cells/mL]); the time at which the carrying capacity was reached, *t-N*_max_ (h); the intrinsic growth rate, *r* (log[cell division/mL/h]); the maximum value of the estimated CO_2_ production rate divided by the estimated cell concentration, *J*_*max*_ (gh^−1^10^−8^cell^−1^); the average cell size at *t-N*_max_, *Size-t-N*_max_(*µ*m); the percentage of living cells at *t-N*_max_, *Viability-t-N*_*max*_ (%); and the percentage of living cells at *t-75, Viability-t-75* (%).
- *Basic enological parameters*, quantified at the end of fermentation: *Residual Sugar* (gL^−1^); *Ethanol* (%vol); the ratio between the amount of metabolized sugar and the amount of released ethanol, *Sugar.Ethanol.Yield* (gL^−1^%vol^−1^); *Acetic acid* (gL^−1^ of H_2_SO_4_); *Total SO*_2_ (mgL^−1^) and *Free SO*_2_ (mgL^−1^).
- *Aromatic traits*, mainly volatile compounds measured at the end of alcoholic fermentation by GC-MS: two higher alcohols (*Phenyl-2-ethanol* and *Hexanol*, mgL^−1^); seven esters (*Phenyl-2-ethanol acetate, Isoamyl acetate, Ethyl-propanoate, Ethyl-butanoate, Ethyl-hexanoate, Ethyl-octanoate* and *Ethyl-decanoate*, mgL^−1^); three medium chain fatty acids (*Hexanoic acid, Octanoic acid* and *Decanoic acid*, mgL^−1^); one thiol *4-methyl-4-mercaptopentan-2-one, X4MMP*(mgL^−1^) and the acetylation rate of higher alcohols, *Acetate ratio*.

For the proteomic analyses samples were harvested at 40 % of CO_2_ release, corresponding to the maximum rate of CO_2_ release. Protein abundances were measured by LC-MS/MS techniques from both shared and proteotypic peptides relying on original Bayesian developments (Blein-Nicolas et al., 2012). A total of 615 proteins were quantified in more than 122 strains × temperature combinations as explained in detail in Blein-Nicolas et al. (2015).

#### 4.1.2 Genetic value of protein abundances and fermentation/life-history traits

We considered the genetic value of protein abundances and fermentation/life-history traits, rather than their measured/computed value. In a previous study, Petrizzelli et al. (2019) decomposed the phenotypic value of a trait at a given temperature, *P*_*T*_, into its genetic, *G*_*T*_, and residual, *ϵ*, contributions:

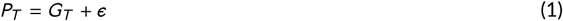

The genetic value, *G*_*T*_, was decomposed in terms of additive and interaction effects, taking into account the structure of the half-diallel design. By including two different species and the parental inbreds in the experimental design, we were able to distinguish between intra- and interspecific additive genetic effects (***A***_***w***_ and ***A***_***b***_, respectively) and decompose the interaction effects into inbreeding (***B***) and intra and interspecific heterosis effects (***H***_***w***_, ***H***_***b***_). Thus, the genetic value of a trait at a given temperature *T* was modeled as follows:

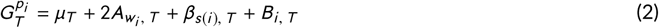

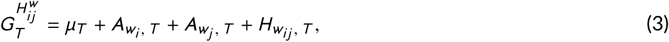

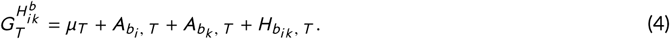

for a parental strain *p*_*i*_ (equation 2), for an intraspecific hybrid 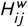 between parents *p*_*i*_ and *p*_*j*_ (eq. 3), and for an interspecific hybrid 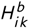 between parents *p*_*i*_ and *p*_*k*_ (eq. 4). *µ* is the overall mean and *β*_*s*(*i*)_ is the deviation from the fixed overall effect for the species:

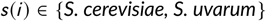

We retrieved the genetic value for all proteomic data. For the fermentation traits, the model did not converge for most ethyl esters (*Ethyl-propanoate, Ethylbutanoate, Ethyl-hexanoate, Ethyl-octanoate* and *Ethyl-decanoate*), and for *Acetate Ratio* and *Acetic acid*. These traits were removed from the final analysis, which in the end included 28 traits.

#### 4.1.3 Protein functional annotation

Cross-referencing MIPS micro-organism protein classification (Ruepp et al., 2004), KEGG pathway classification (Kanehisa and Goto, 2000; Kanehisa et al., 2016, 2017) and Saccharomyces Genome database (Cherry et al., 2012b) information, we attributed a single functional category to each protein.

The first two hierarchical levels of MIPS functional annotation were taken into account to assign proteins to 34 different categories. All secondary levels were used for the *01.metabolism, 02.energy* and *10.cell cycle and DNA processing* categories, resulting in 20 different functional categories. The *11.transcription* category was subdivided into the *transcription* sub-group (*11.06 and 11.02*) and into the *RNA processing* sub-group (*11.04*). Similarly, the *12.protein synthesis* category was split into the *ribosomal proteins* (*12.01*) and *translation* (*12.04, 12.07, 12.10*) sub-groups, and the *20.transport* category was split into the *vacuolar transport* (*20.09*) and *transport* (*20.01, 20.03*) sub-groups.

By contrast, the first hierarchical category was used for *14.protein fate, 30.signal transduction, 32.detoxif,cation, 34.homeostasis, 40.cell growth and death, 42.cytoskeleton* In addition, *16.binding function* and *18.02.regulation* category were fused into *16.binding*, and *32.transposon movement* was fused with *10.01.DNA processing*. Finally, *41.mating* and *43.budding* were included in the *10.03.cell cycle* category.

#### 4.1.4 DynamosYeast model

We used the DynamoYeast model, which is a previously developed Constraint-Based Model of central carbon metabolism in *S. cerevisiae* (Celton et al., 2012). The main metabolic pathways included in this model are upper and lower glycolysis, the pentose phosphate pathway (PPP), glycerol synthesis, ethanol synthesis and the reductive and oxidative branches of the tricarboxylic acid (TCA) cycle. This model consists of 60 metabolites and 70 reactions, including one input flux, the uptake of glucose, and 10 output fluxes (Fig 1), taking place in the cytosol, mitochondria or in the extracellular medium.

The range of variation of the fluxes was fixed to allow for alcoholic fermentation. The following reactions were considered to be irreversible with ***v***^inf^ = 0: *Oaa_Mal* (malate dehydrogenase), *Mal_Fum* (fumarase), *Fum_Succ* (fumarate reductase), and their respective mitochondrial counterparts *Oaa_Mal_m, Mal_Fum_m* and *Fum_Succ_m*, and *Oaa_Cit_m* (mitochondrial citrate synthase) (fluxes are denoted through the abbreviation of the substrate and the product connected by “_”, followed by the enzyme name in parentheses). The fructose reaction was not included in the model, and *Glu_Akg_m* (mitochondrial glutamate dehydrogenase), as well as *Aceto_But* (butanediol dehydrogenase) were set to zero. In all, there were 16 reversible and 52 irreversible reactions.

Following the conventions implemented by many genome-scale metabolic models, many reactions of the DynamoYeast model of central carbon metabolism in *S. cerevisiae* are associated with genes and proteins via gene-protein-reaction (GPR) associations (Thiele and Palsson, 2010).

In general, there can be a many-to-many mapping of genes to reactions; for example, one reaction can be linked to proteins (*P* 1 and *P* 2) or *P* 3. The first Boolean AND relationship means that the reaction is catalyzed by a complex of two gene products. Since the maximum of the complex is given by the minimum of its components, the weighting of the complex is defined as: *P* 1 AND *P* 2 = min(*P* 1, *P* 2). The OR relationship allows for alternative catalysts of the reaction. Thus, total capacity is given by the sum of its components: (*P* 1 AND *P* 2) OR *P* 3 = min(*P* 1, *P* 2)+ *P* 3 (Lee et al., 2012). Following these rules for each of the 11 yeast strains and the 55 hybrids at both temperatures, we estimated the protein abundances associated with the reactions of the DynamoYeast model, resulting in 33 reaction weightings out of 70.

### 4.2 Methods

#### 4.2.1 Constraint-based modeling of metabolic networks

Metabolic networks can be described in terms of the relationship between *M* metabolites, ***m***, and *N* reactions, ***v***, at a given time *t*:

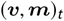

Their topology can be expressed through the *M* × *N* stoichiometric matrix ***S***, in which rows correspond to the stoichiometric coefficients of the corresponding metabolites in all reactions.

Under mass-balance assumptions and thermodynamic bounds of reaction rates, the dynamics of the network are governed by the linear system of constraints and inequalities:

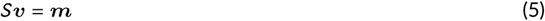

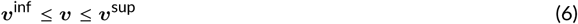

where ***m*** ∈ ℝ^*M*^ is the vector of the *M* metabolite input/output rates, ***v*** ∈ ℝ^*N*^ is the set of *N* reactions, and ***v***^inf^, ***v***^sup^ are the extremes of variation of the set of fluxes. Under a steady state assumption, ***m*** = 0 and the feasible solution space is expressed as:

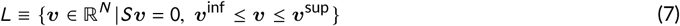

In general, *N* is larger than *M* and the solution space *L* has infinite cardinality.

#### 4.2.2 Characterization of the feasible solution space

We characterized the feasible solution space *L* through the posterior probability of flux values obtained by the Expectation Propagation (EP) algorithm described in Braunstein et al. (2017).

Instead of exploring *L* by sampling, as classical methods do, Braunstein et al. (2017) combined statistical physics and Bayesian approaches to infer the joint distribution of metabolic fluxes. To do so, given a set of metabolite input/output rates, ***m***, they encoded the stoichiometric constraints within the likelihood posterior probability, defining a Boltzmann-like distribution with an energetic quadratic function

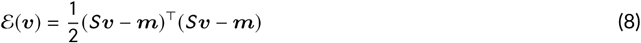

while the inequality constrains were encoded in the prior probability of fluxes. Using Bayes theorem, this method provides a model for the posterior density of flux distribution.

Therefore, each point ***v*** in *L* follows the truncated multivariate normal distribution

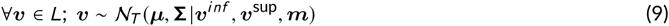

where ***µ*** is the vector of the mean posterior values of fluxes and **Σ** the posterior variance-covariance matrix of fluxes estimated with the EP algorithm.

For each set of metabolic fluxes ***v***, the posterior probability of observing ***v*** can be calculated at follows:

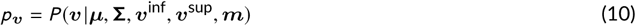

Different values of extremes of variation can be used to model a particular process, for example for modeling reactions known to be irreversible in a specific context, *i. e*.

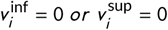

or for introducing experimental data constraints, *i. e*.

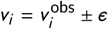

for the *i* -th reaction.

Given that ***µ*** and **Σ** depend on the imposed range (***v***^inf^, ***v***^sup^) of internal and exchange fluxes, metabolic fluxes will take on particular values with probabilities that will depend on *a priori* knowledge and on the chosen metabolic process.

The algorithm implemented by Braunstein et al. (2017) was translated into *R* code. Extraction of the stoichiometric matrix from the DynamosYeast model was performed with the *sybil* package in R (Gelius-Dietrich et al., 2013).

#### 4.2.3 Prediction of metabolic fluxes from proteomic data

In living systems, most metabolic reactions are catalyzed by enzymes, and quantitative proteomic data retain information about enzyme abundances. Therefore, the metabolism of a cell at a given time is characterized by the set of fluxes, metabolites and protein abundances

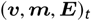

where ***E*** = (*E*_1_, *E*_2_, …, *E*_*N*_), and *E*_*i*_ is the abundance of enzyme *i* associated with the reaction flux ***v***_*i*_. Indeed, even though reaction rates are not directly proportional to enzyme abundances, a degree of covariation between protein abundance and flux reaction rate can be expected at the scale of the metabolic network. This can be used to infer intracellular metabolic fluxes with reasonable accuracy (Lee et al., 2012).

Among all possible solutions in *L*, we chose the one that minimized the objective function:

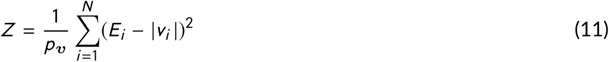

*i.e*. the Euclidean distance between the observed protein abundances ***E***_*obs*_ and the associated fluxes, weighted by *p*_***v***_, the posterior probability of observing the set of metabolic fluxes ***v***.

The properties of the truncated multivariate normal distribution ensure that the solution of the objective function is unique and no sophisticated algorithm is needed to find this solution. For each set of observations ***E***_obs_, we sampled *N*_*s*_ points of the feasible solution space. Therefore, ∀*k* ∈ {1, 2, 3 … *N*_*s*_ }, we obtained ***v***^*k*^ ∈ *L* and 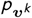. We calculated *Z* ^(*k*)^ and selected the set of flux values, ***v***^predicted^, for which *Z* ^(*k*)^ was the minimum.

In practice, it is never possible to associate each reaction of the metabolic network with a protein abundance. First, quantitative proteomics is not exhaustive. Second, reactions in a metabolic model are not always associated with an enzyme. Assuming a steady state condition and introducing information about protein abundances and measured external metabolic fluxes allows us to describe the system as:

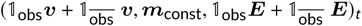

where 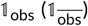 is an indicator vector: its component-wise value is equal to 1 if the associated flux/protein component is observed (unobserved), otherwise it is equal to 0. Taking this into account, we reformulated the problem as follows:

- Observed fluxes were introduced as additional constraints with

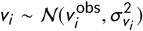

where 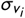 was set to a small value.
- The objective function was calculated only on the subset of observed enzyme abundances:

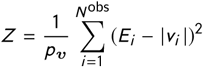

Predictions of metabolic fluxes were performed by coupling the DynamoYeast model to our experimental data (protein abundances and CO_2_ reaction rates, the only measured flux in our study). We constrained the solution space *L* by considering the maximum CO_2_ release rate, measured at the same time point as the one used in the proteomics analyses (Blein-Nicolas et al., 2015). For each strain × temperature combination a unique solution was obtained by minimizing the objective function, defined in equation 11, from the set of observations.

#### 4.2.4 Testing the prediction algorithm

The prediction algorithm is based on the assumption that fluxes and enzyme abundances covary. Indeed, any reaction rate can be expressed as a more or less complex function of enzyme abundances, kinetic constants and metabolite concentrations (Fell and Cornish-Bowden, 1997):

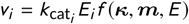

where *k*_*cat*_ is the catalytic constant, *κ* is a set of other kinetic constants, *E* is the set of enzymes abundances other than enzyme *i*. The *f* function can be more or less complex depending on the mode of regulation.

To test the accuracy of the prediction of metabolic fluxes from protein abundance data, we used the feasible solution space of the DynamoYeast model and three different functions relating reaction rates to enzyme abundances. Specifically, we reversed the relationship, expressing protein abundance as a function of the reaction rate using a simplified formalism derived from the Metabolic Control Theory (Kacser and Burns, 1973; Heinrich and Rapoport, 1974)

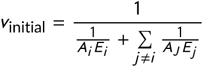

where the *A*^′^_*j*_ *s* are positive or negative constant terms. Given that enzyme concentrations cannot be negative, and taking ∀*j, A*_*j*_ = ±1, we obtain the hyperbolic relation:

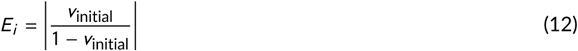

We also tested the predictions under the assumption that the relationship between protein abundances and flux reaction rates was linear:

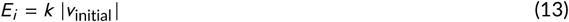

*k* being an uniform random number *k* ∼ 𝒰 (0.1, 3)

Finally, we considered the case where protein abundances and flux reaction rates are linked by a sigmoidal function (Nijhout et al., 2003), which we approximated with a Hill function:

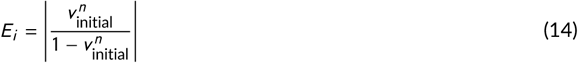

where *n* is the Hill coefficient, sampled in the set Ω = {2, 3, 4, 5}.

For each simulation, we sampled an initial set of fluxes ***v***_initial_ ∈ *L*. We estimated the complete set of enzymatic protein abundances, ***E***_initial_ using equations (12, 13 or 14). Then, we minimized the objective function *Z* to predict the set of fluxes ***v***^predicted^ that best fit enzyme abundances. Prediction accuracy was measured as the correlation coefficient between ***v***^predicted^ and ***v***_initial_. Computer simulations were performed to test the influence of two main parameters: (*i*) the number of sampled points *N*_*s*_; (*i i*) the number of quantified proteins, *N* ^obs^, included in the minimization process.

We assumed a steady state condition (***m*** = 0) and sampled *N*_*s*_ points from the solution space of the multivariate posterior joint distribution of fluxes obtained using the EP algorithm (Braunstein et al., 2017). We drew an additional point in solution space *L*, ***v***_initial_, and we calculated protein concentrations from the inverse problem. We retained the set of fluxes, ***v***^predicted^ for which *Z* was minimum. The numbers *N* ^obs^ and *N*_*s*_ were let to vary (*N*_*s*_ ∈ {10^2^, 10^3^, 10^4^, 10^5^, 10^6^ } and *N* ^obs^ ∈ {1, 2, 3 … }).

In terms of computational time, it would be expensive to consider all the possible combinations of observed enzymatic proteins associated with the metabolic model that can be included in equation 11 (there are *N* ^obs^(1 + (*N* ^obs^ − 1) + (*N* ^obs^ − 1)(*N* ^obs^ − 2) + … + (*N* ^obs^ − 1)!) combinations). Therefore, for a given *N*_*s*_, our strategy was to randomly choose one-by-one a protein to be included in the computation of the *Z* function, and therefore for the prediction of metabolic fluxes ***v***^predicted^.

We randomly choose one reaction, *v*_1_, from the complete set of reactions in the model, and we minimized

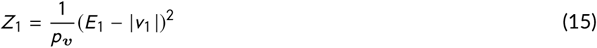

to select one solution 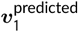 over the *N*_*s*_ possible solutions of *L*. At the next iteration, we randomly chose an additional flux *v*_2_ and its associated protein abundance *E*_2_, and we minimized

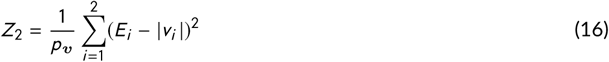

to predict 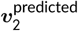. This procedure was performed until the complete set of reactions was selected. In total, simulations were run a thousand times for different values of *N*_*s*_ and *N* ^obs^.

### 4.3 Statistical Analysis

In order to study the main features characterizing fermentation and life-history traits in the HeterosYeast dataset, we analyzed the variation components of data from three different levels of cellular organization: protein abundances **E**, metabolic fluxes **V** and fermentation/life-history traits **T**:

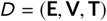

The total number of observations was 127 strain × temperature combinations (66 strains, 2 temperatures, minus 5 missing data due to the poor fermenting abilities of some strains). The whole dataset consisted of 615 protein abundances, 70 metabolic fluxes and 28 fermentation and life-history traits. Two types of analyses were performed using different multivariate approaches: analyses at a single phenotypic level and analyses integrating the different levels.

We ran Principal Component Analyses (PCA) to identify the most important sources of variation within the datasets and the similarities/differences between the different phenotypic levels. We included prior knowledge regarding the yeast species in order to perform a supervised sparse Partial Least Squares Discriminant Analysis (sPLS-DA) to extract and combine discriminating features that best separate the different groups. The number of selected features was tuned using 3-fold cross-validation repeated 1,000 times.

In addition, the three levels of cellular organization were integrated in an unsupervised framework by performing a regularized Canonical Correlation Analysis (rCCA), using the *mixomics* package in R (Lê Cao et al., 2009; Rohart et al., 2017). First, we searched for the key features that maximized the correlation between metabolic fluxes and fermentation traits. Second, we looked for groups of proteins that maximized the correlation with the most integrated traits (regularization parameters were tuned using a leave-one-out cross-validation procedure on a 1000 × 1000 grid between 0.0001 to 1). Finally, Pearson’s chi-square test of enrichment was computed on protein functional category frequencies taking as prior probability the expected category frequency found in the MIPS database.

Since the correlation matrix between traits and fluxes was clearly structured, we computed the matrix of Euclidean distance between traits, based on their correlation with metabolic fluxes, and clustered traits using the *hcl ust* package in *R*. This procedure allowed us to define five groups of traits that showed similar correlation patterns with fluxes of the central carbon metabolism. Finally, we stored the linear correlation coefficients between proteins (P = 615) and traits (T = 28) in a (*T* × *P*) and ran a Linear Discriminant Analysis to search for proteins that best discriminate between groups of traits, considering traits as individuals. A functional analysis of the proteins that best correlated with LDA axes was performed using the 34 protein functional categories defined above.

## Data availability

The datasets and computer codes produced in this study are available at *f,gshare*, DOI:10.6084/m9.figshare.10266332. The folder includes:

- The *R* code (translated from matlab) of the Expectation Propagation algorithm implemented in Braunstein et al. (2017);
- The genetic values of the protein abundances in the HeterosYeast dataset retrieved from Petrizzelli et al. (2019);
- The genetic values of the fermentation and life-history traits in the HeterosYeast dataset retrieved from Petrizzelli et al. (2019);
- The predicted values of the metabolic fluxes of the central carbon metabolism (Celton et al., 2012) for each strain × temperature combination in the HeterosYeast dataset.

## Acknowledgements

This work was funded with a public Ph.D. grant from the French National Research Agency (ANR) as part of the Investissement d’Avenir program, through the Initiative Doctorale Interdisciplinaire (IDI) 2015 project funded by the Initiative d’Excellence (IDEX) Paris-Saclay, ANR-11-IDEX-0003-02.

## Author contribution

Christine Dillmann and Dominique de Vienne proposed the project and with Marianyela Petrizzelli developed and implemented the methodology. Thibault Nidelet calibrated the metabolic model of fluxes of the central carbon metabolism to simulate alcoholic fermentation. MP translated the Expectation Propagation algorithm in R and with CD performed all the statistical analyses. All authors wrote, edited and approved the manuscript.

## S1 SUPPLEMENTARY METHODS

### S1.1 Sampling the solution space

Let *L* denote the solution space of eq. 5 with constraints (eq. 6). Our aim is to sample random elements in the convex set *L* in order to characterize it by means of the posterior joint distribution of fluxes. This can be achieved using classical methods, such as the well-known **Hit and Run** algorithm (Meersche et al., 2009). Braunstein et al. (2017) turned to map the original problem of sampling the feasible solution space *L* into an inference problem of the joint distribution of metabolic fluxes, letting the linear and inequality constraints to be encoded within the likelihoods and prior probabilities, which via Bayes theorem provides a model for the posterior distribution density of the flux.

We compared the posterior density distribution obtained by Hit and Run (HR) sampling with the Expectation Propagation algorithm (EP). We ran HR with a burn-in length equal to 10^6^ and a jump of 0.5, for a number of 10^6^ to 10^7^ iterations, and the EP algorithm with a high *β* parameter (Boltzmann inverse temperature parameter). Fig. S1 hows the solution space sampled by HR (histograms) and the EP estimate (red curve). Fig. S2 shows the Pearson correlation coefficients between variances and means estimated with EP and HR for different numbers of iterations. As can be seen, the Pearson correlation increases as the number of HR samples increases. Assuming that HR samples the true distribution of fluxes, means are well predicted by the EP algorithm, although variances are underestimated.

We further investigated whether the EP algorithm predicted well the variance-covariance matrix of the DynamoYeast fluxes. Fig. S3 shows the relationship between 8 pairwise fluxes chosen at random, and the correlation ellipses (red curve) computed by the EP algorithm. As can be seen, the EP algorithm predicts well the variance-covariance matrix of fluxes, satisfying eq. 5, based on HR predictions.

### S2 SUPPLEMENTARY FIGURES

**FIGURE S1.**
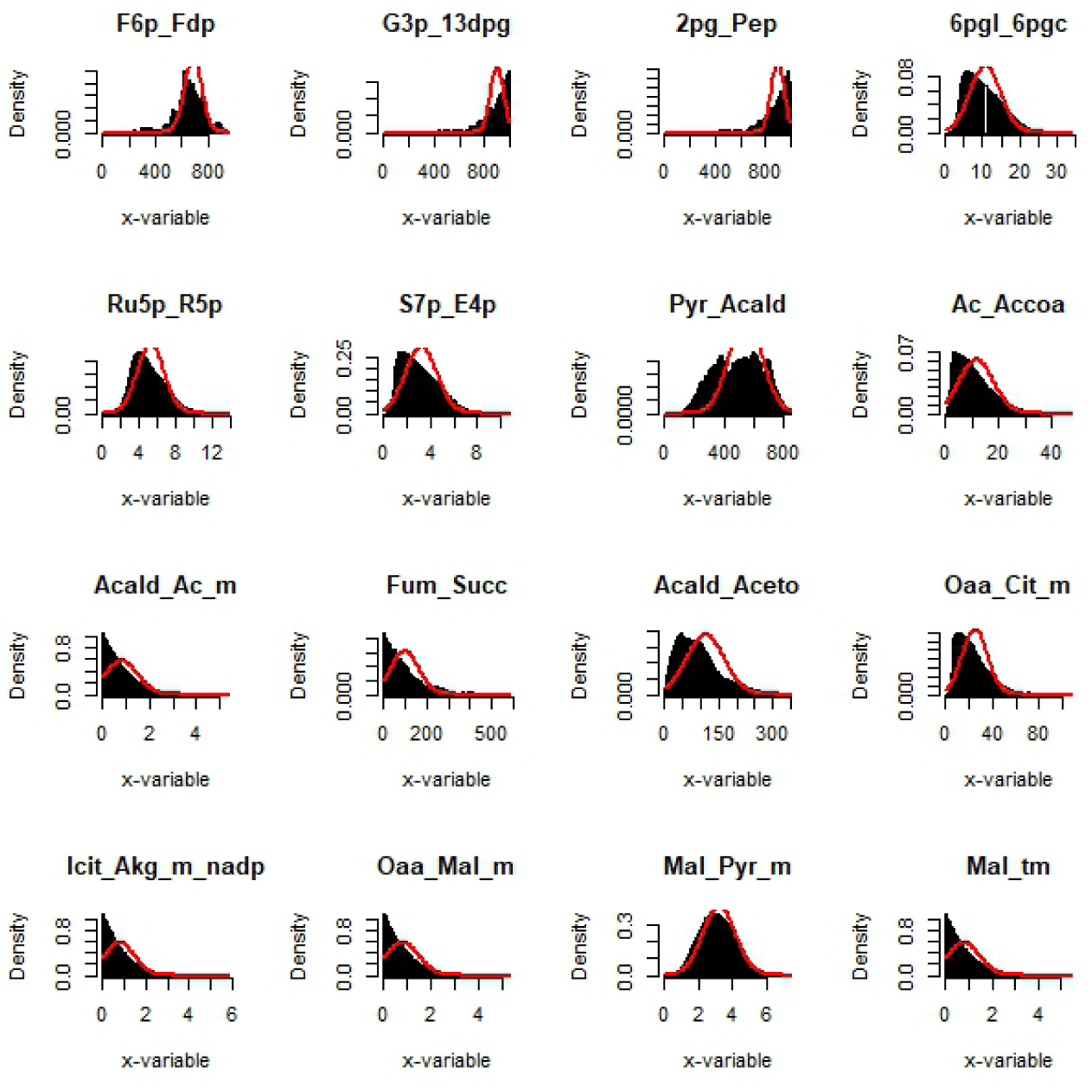
Marginal probability densities of sixteen randomly chosen fluxes of carbon metabolism in yeast. The histograms represent the HR result for *T* ∼ 10^7^ sampling points. The red line is the result of the EP estimate.

**FIGURE S2.**
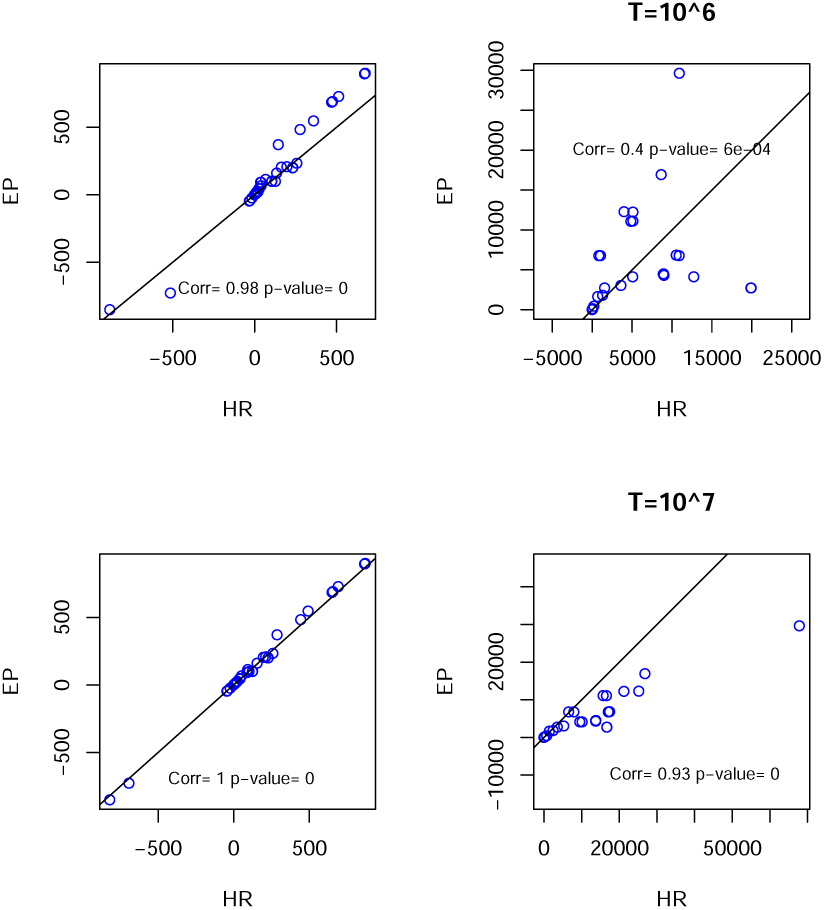
Comparison of the results of HR versus EP. The plots on the left are scatter plots of the means and on the right variances of the approximated marginals computed via EP against the ones estimated via HR for an increasing number of explored configurations *T*, top *T* ∼ 10^6^, bottom *T* ∼ 10^7^.

**FIGURE S3.**
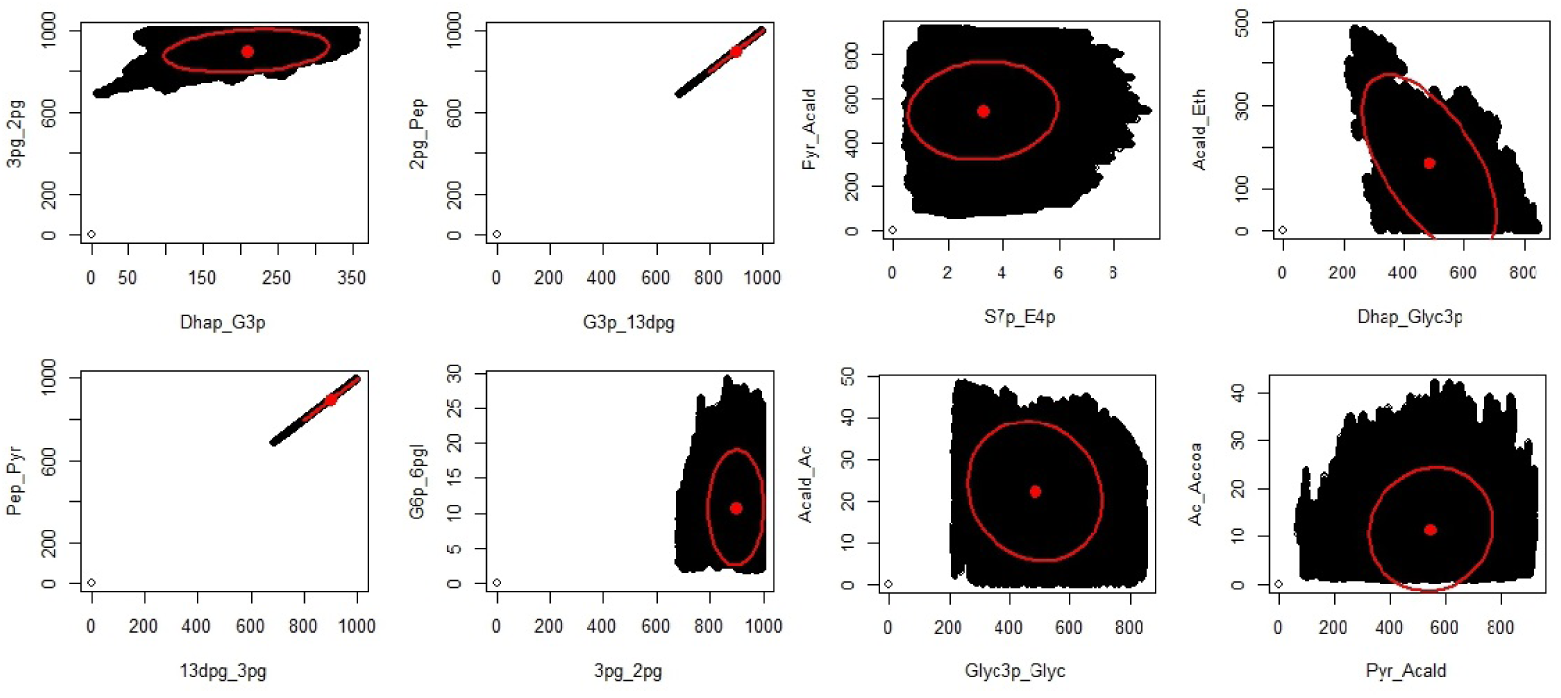
Comparison of the results of HR versus EP. The plot shows the relationship between 8 pairwise fluxes. Correlation ellipses computed by the EP algorithm are drawn in red. Dot points represent the mean value of fluxes computed with EP. For HR samples, *T* ∼ 5 ∗ 10^6^.

**FIGURE S4.**
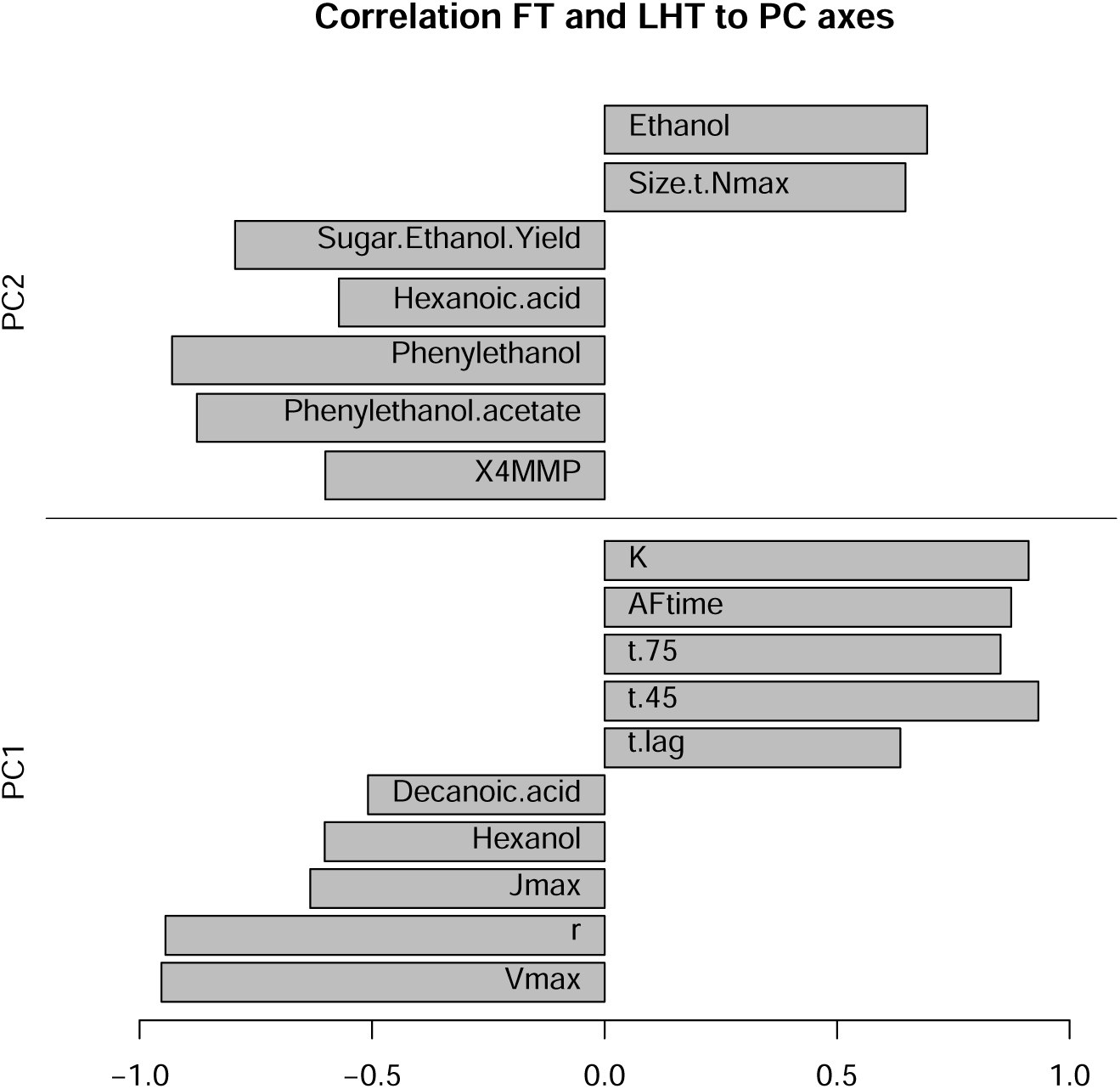
Correlation between fermentation and life-history traits and the first two axes of the Principal Component Analysis. The figure shows traits for which the correlation was >0.5 or <-0.5 (p-value < 0.05). The first axis is negatively correlated with growth rate (*r*), CO_2_ fluxes (*J*_max_ and *V*_max_), *Hexanol* and *Decanoic acid* and positively correlated with carrying capacity (*K*) and fermentation times (*AFtime, t-lag, t-75, t-45*). The second axis is positively correlated with cell size (*Size-t-N*_max_) and *Ethanol* at the end of fermentation, and negatively correlated with aroma production at the end of fermentation and *Sugar.Ethanol.Yield*.

**FIGURE S5.**
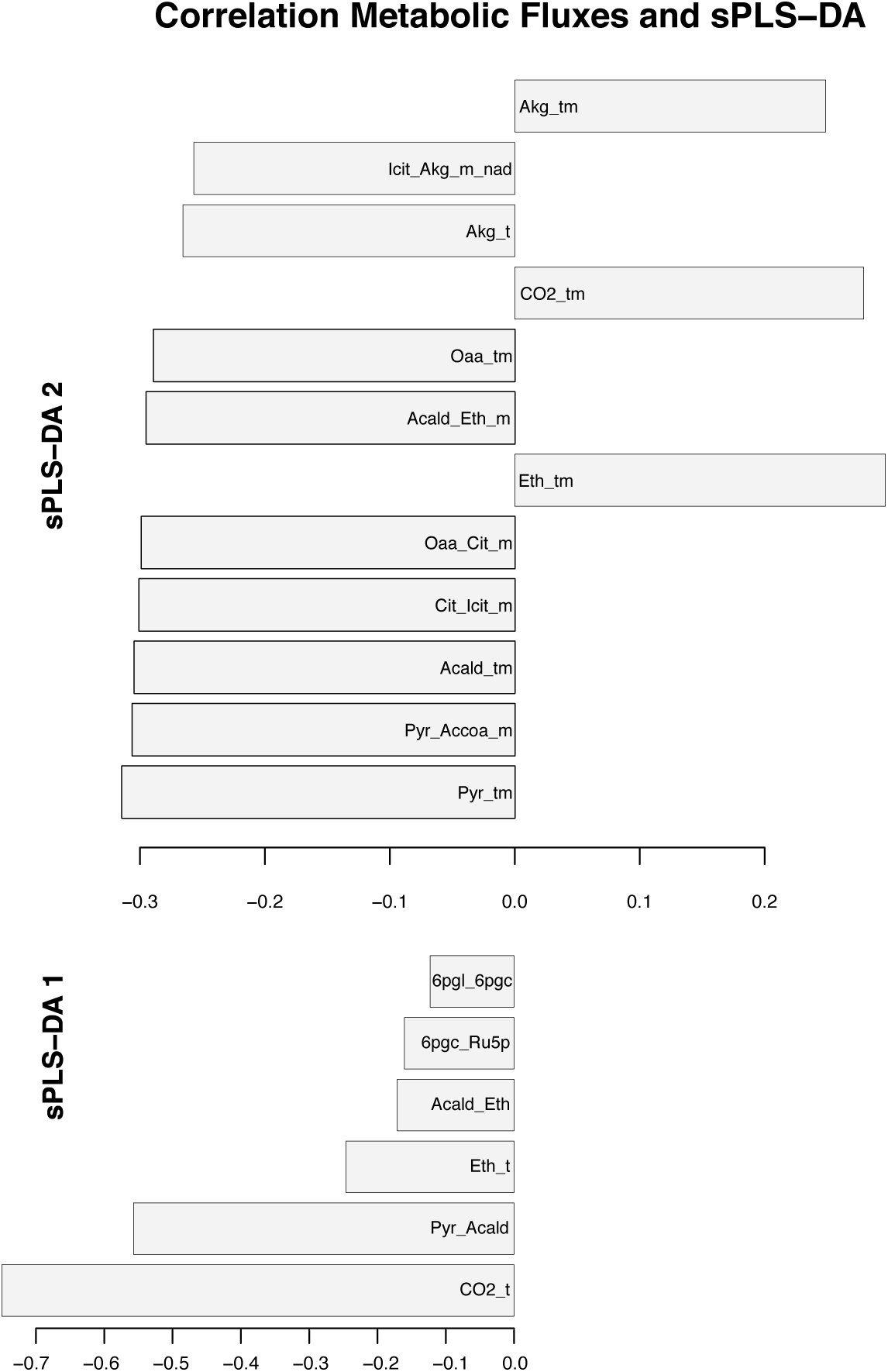
Correlation between metabolic fluxes and the first two axes of the sparse Partial Least Square Discriminant Analysis. The CO_2_, pyruvate decarboxylase, ethanol, alcohol dehydrogenase, phosphogluconolactonase and phosphogluconate dehydrogenase fluxes contributed to the first axis of the sPLS-DA, and were all negatively correlated with it. The second axis was negatively correlated with the mitochondrial acetyl-CoA formation, mitochondrial citrate synthase, mitochondrial aconitate hydratase, mitochondrial isocitrate dehydrogenase (NAD+) and mitochondrial transport fluxes of pyruvate, oxaloacetate and acetaldehyde fluxes, while positively correlated with the mitochondrial transport of 2-oxodicarboylate, ethanol and CO_2_ fluxes.

## References

Albertin, W., Marullo, P., Bely, M., Aigle, M., Bourgais, A., Langella, O., Balliau, T., Chevret, D., Valot, B., Silva, T. d., Dillmann, C., Vienne, D. d. and Sicard, D. (2013a) Linking Post-Translational Modifications and Variation of Phenotypic Traits. Molecular & Cellular Proteomics, 12, 720–735. URL: http://www.mcponline.org/content/12/3/720.

Albertin, W., da Silva, T., Rigoulet, M., Salin, B., Masneuf-Pomarede, I., de Vienne, D., Sicard, D., Bely, M. and Marullo, P. (2013b) The mitochondrial genome impacts respiration but not fermentation in interspecific saccharomyces hybrids. PLOS ONE, 8, 1–14.

Antoniewicz, M. (2015) Methods and advances in metabolic flux analysis: a mini-review. J Ind Microbiol Biotechnol., 42, 317–25.

Belouah, I., Nazaret, C., Pétriacq, P., Prigent, S., Bénard, C., Mengin, V., Blein-Nicolas, M., Denton, A. K., Balliau, T., Augé, S., Bouchez, O., Mazat, J.-P., Stitt, M., Usadel, B., Zivy, M., Beauvoit, B., Gibon, Y. and Colombié, S. (2019) Modeling protein destiny in developing fruit. Plant Physiology, pp.00086.2019. URL: http://www.plantphysiol.org/content/early/2019/04/29/pp.19.00086.

Blein-Nicolas, M., Albertin, W., da Silva, T., Valot, B., Balliau, T., Masneuf-Pomarede, I., Bely, M., Marullo, P., Sicard, D., Dillmann, C., de Vienne, D. and Zivy, M. (2015) A systems approach to elucidate heterosis of protein abundances in yeast. Mol Cell Proteomics, 14, 2056–71.

Blein-Nicolas, M., Albertin, W., Valot, B., Marullo, P., Sicard, D., Giraud, C., Huet, S., Bourgais, A., Dillmann, C., de Vienne, D. and Zivy, M. (2013) Yeast proteome variations reveal different adaptive responses to grape must fermentation. Molecular Biology and Evolution, 30, 1368. URL: +http://dx.doi.org/10.1093/molbev/mst050.

Blein-Nicolas, M., Xu, H., de Vienne, D., Giraud, C., Huet, S. and Zivy, M. (2012) Including shared peptides for estimating protein abundances: A significant improvement for quantitative proteomics. PROTEOMICS, 12, 2797–2801. URL: http://dx.doi.org/10.1002/pmic.201100660.

Braunstein, A., Muntoni, A. P. and Pagnani, A. (2017) An analytic approximation of the feasible space of metabolic networks. Nature Communications, 8, 14915. URL: http://www.nature.com/doifinder/10.1038/ncomms14915.

Bélisle, C. J. P., Romeijn, H. E. and Smith, R. L. (1993) Hit-and-Run Algorithms for Generating Multivariate Distributions. Mathematics of Operations Research, 18, 255–266. URL: https://www.jstor.org/stable/3690278.

Caspi, R., Altman, T., Billington, R., Dreher, K., Foerster, H., Fulcher, C., Holland, T., Keseler, I., Kothari, A., Kubo, A., Krummenacker, M., Latendresse, M., Mueller, L., Ong, Q., Paley, S., Subhraveti, P., Weaver, D., Weerasinghe, D., Zhang, P. and Karp, P. (2014) The metacyc database of metabolic pathways and enzymes and the biocyc collection of pathway/genome databases. Nucleic Acids Res., 42(Database issue), D459–71.

Celton, M., Goelzer, A., Camarasa, C., Fromion, V. and Dequin, S. (2012) A constraint-based model analysis of the metabolic consequences of increased NADPH oxidation in Saccharomyces cerevisiae. Metabolic Engineering, 14, 366 – 379.

Cherry, J., Hong, E., Amundsen, C., Balakrishnan, R., Binkley, G., Chan, E., Christie, K., Costanzo, M., Dwight, S., Engel, S., Fisk, D., Hirschman, J., Hitz, B., Karra, K., Krieger, C., Miyasato, S., Nash, R., Park, J., Skrzypek, M., Simison, M., Weng, S. and Wong, E. (2012a) Saccharomyces genome database: the genomics resource of budding yeast. Nucleic Acids Res., 40(Database issue), D700–5.

Cherry, J. M., Hong, E. L., Amundsen, C., Balakrishnan, R., Binkley, G., Chan, E. T., Christie, K. R., Costanzo, M. C., Dwight, S. S., Engel, S. R., Fisk, D. G., Hirschman, J. E., Hitz, B. C., Karra, K., Krieger, C. J., Miyasato, S. R., Nash, R. S., Park, J., Skrzypek, M. S., Simison, M., Weng, S. and Wong, E. D. (2012b) Saccharomyces Genome Database: the genomics resource of budding yeast. Nucleic Acids Research, 40, D700–705.

Collot, D., Nidelet, T., Ramsayer, J., Martin, O. C., Méléard, S., Dillmann, C., Sicard, D. and Legrand, J. (2018) Feedback between environment and traits under selection in a seasonal environment: consequences for experimental evolution. Proceedings of the Royal Society B: Biological Sciences, 285, 20180284. URL: http://rspb.royalsocietypublishing.org/lookup/doi/10.1098/rspb.2018.0284.

Edwards, J., Ibarra, R. and Palsson, B. (2001) In silico predictions of escherichia coli metabolic capabilities are consistent with experimental data. Nature Biotechnology, 19, 125–130.

Fell, D. and Cornish-Bowden, A. (1997) Understanding the control of metabolism, vol. 2. Portland press London.

Fell, D. and Small, J. (1986) Fat synthesis in adipose tissue. an examination of stoichiometric constraints. Biochem J., 238, 781–786.

Fisher, R. A. (1930) The Genetical Theory of Natural Selection. Clarendon Press, Oxford.

Gelius-Dietrich, G., Fritzemeier, C. J., Desouki, A. A. and Lercher, M. J. (2013) sybil – efficient constraint-based modelling in r. BMC Systems Biology, 7, 125. URL: http://www.biomedcentral.com/1752-0509/7/125.

Gould, S. and Lewontin, R. (1979) The spandrels of san marco and the panglossian paradigm: a critique of the adaptationist programme. Proc. R. Soc. Lond. B, 295, 581–598.

Heinrich, R. and Rapoport, T. A. (1974) A linear steady-state treatment of enzymatic chains. general properties, control and effector strength. 42, 89–95.

Kacser, H. and Burns, J. A. (1973) The control of flux. 27, 65–104.

Kanehisa, M., Furumichi, M., Tanabe, M., Sato, Y. and Morishima, K. (2017) KEGG: new perspectives on genomes, pathways, diseases and drugs. Nucleic Acids Research, 45, D353–D361.

Kanehisa, M. and Goto, S. (2000) KEGG: kyoto encyclopedia of genes and genomes. Nucleic Acids Research, 28, 27–30.

Kanehisa, M., Sato, Y., Kawashima, M., Furumichi, M. and Tanabe, M. (2016) KEGG as a reference resource for gene and protein annotation. Nucleic Acids Research, 44, D457–462.

Lee, D., Smallbone, K., Dunn, W. B., Murabito, E., Winder, C. L., Kell, D. B., Mendes, P. and Swainston, N. (2012) Improving metabolic flux predictions using absolute gene expression data. BMC Systems Biology, 6, 73. URL: https://doi.org/10.1186/1752-0509-6-73.

Lê Cao, K.-A., González, I. and Déjean, S. (2009) integrOmics: an R package to unravel relationships between two omics datasets. Bioinformatics (Oxford, England), 25, 2855–2856.

Meersche, K. V. d., Soetaert, K. and Oevelen, D. V. (2009) xsample(): An *R* Function for Sampling Linear Inverse Problems. Journal of Statistical Software, 30. URL: http://www.jstatsoft.org/v30/c01/.

Nicholson, J. and Lindon, J. (2008) Systems biology: Metabonomics. Nature, 455, 1054–6.

Nidelet, T., Brial, P., Camarasa, C. and Dequin, S. (2016) Diversity of flux distribution in central carbon metabolism of S. cerevisiae strains from diverse environments. Microbial Cell Factories, 15. URL: http://microbialcellfactories.biomedcentral.com/articles/10.1186/s12934-016-0456-0.

Nijhout, H. F., Berg, A. M. and Gibson, W. T. (2003) A mechanistic study of evolvability using the mitogen-activated protein kinase cascade. Evolution & Development, 5, 281–294. URL: https://onlinelibrary.wiley.com/doi/abs/10.1046/j.1525-142X.2003.03035.x.

Palsson, B. O. (2015) Systems Biology, Constraint-based Reconstruction and Analysis. Cambridge University Press.

Petrizzelli, M., Vienne, D. d. and Dillmann, C. (2019) Decoupling the Variances of Heterosis and Inbreeding Effects Is Evidenced in Yeast’s Life-History and Proteomic Traits. Genetics, 211, 741–756. URL: https://www.genetics.org/content/211/2/741.

Pianka, E. R. (1970) On r- and K-Selection. The American Naturalist, 104, 592–597. URL: https://www.journals.uchicago.edu/doi/10.1086/282697.

Riekeberg, E. and Powers, R. (2017) New frontiers in metabolomics: from measurement to insight. F1000Research, 6, 1148.

Rohart, F., Gautier, B., Singh, A. and Cao, K.-A. L. (2017) mixOmics: An R package for ‘omics feature selection and multiple data integration. PLOS Computational Biology, 13, e1005752. URL: https://journals.plos.org/ploscompbiol/article?id=10.1371/journal.pcbi.1005752.

Ruepp, A., Zollner, A., Maier, D., Albermann, K., Hani, J., Mokrejs, M., Tetko, I., Güldener, U., Mannhaupt, G., Münsterkötter, M. and Mewes, H. W. (2004) The FunCat, a functional annotation scheme for systematic classification of proteins from whole genomes. Nucleic Acids Research, 32, 5539–5545.

Sabarly, V., Aubron, C., Glodt, J., Balliau, T., Langella, O., Chevret, D., Rigal, O., Bourgais, A., Picard, B., Vienne, D. d., Denamur, E., Bouvet, O. and Dillmann, C. (2016) Interactions between genotype and environment drive the metabolic phenotype within Escherichia coli isolates. Environmental Microbiology, 18, 100–117. URL: https://onlinelibrary.wiley.com/doi/abs/10.1111/1462-2920.12855.

da Silva, T., Albertin, W., Dillmann, C., Bely, M., la Guerche, S., Giraud, C., Huet, S., Sicard, D., Masneuf-Pomarede, I., de Vienne, D. and Marullo, P. (2015) Hybridization within saccharomyces genus results in homoeostasis and phenotypic novelty in winemaking conditions. PLOS ONE, 10, 1–24. URL: http://dx.doi.org/10.1371%2Fjournal.pone.0123834.

Spor, A., Nidelet, T., Simon, J., Bourgais, A., de Vienne, D. and Sicard, D. (2009) Niche-driven evolution of metabolic and life-history strategies in natural and domesticated populations of Saccharomyces cerevisiae. BMC Evolutionary Biology, 9, 296. URL: https://doi.org/10.1186/1471-2148-9-296.

Spor, A., Wang, S., Dillmann, C., Vienne, D. d. and Sicard, D. (2008) “Ant” and “Grasshopper” Life-History Strategies in Saccharomyces cerevisiae. PLOS ONE, 3, e1579. URL: https://journals.plos.org/plosone/article?id=10.1371/journal.pone.0001579.

Stearns, S. (1992) The evolution of life histories. Oxford University Press.

Thiele, I. and Palsson, B. O. (2010) A protocol for generating a high-quality genome-scale metabolic reconstruction. Nature protocols, 5, 93–121. URL: https://www.ncbi.nlm.nih.gov/pmc/articles/PMC3125167/.

Töpfer, N., Kleessen, S. and Nikoloski, Z. (2015) Integration of metabolomics data into metabolic networks. Frontiers in plant science, 6, 49.

Wagner, G. P. and Zhang, J. (2011) The pleiotropic structure of the genotype-phenotype map: the evolvability of complex organisms. Nat Rev Genet., 12, 204–13.

Watson, M. (1984) Metabolic maps for the apple ii. Biochemical Society Transactions, 12, 1093–1094.

